# Normothermic human kidney preservation drives iron accumulation and ferroptosis

**DOI:** 10.1101/2025.02.24.639914

**Authors:** Marlon J.A. de Haan, Marleen E. Jacobs, Annemarie M.A. de Graaf, Roan H. van Scheppingen, Rico J.E. Derks, Dorottya K. de Vries, Jesper Kers, Ian P.J. Alwayn, Cees van Kooten, Elena Sánchez-López, Martin Giera, Marten A. Engelse, Ton J. Rabelink

## Abstract

Ex vivo normothermic machine perfusion has been proposed to protect deceased donor organs, promoting metabolic recovery and allowing quality assessment. However, its benefits for preserving deceased donor kidneys remain ambiguous. We postulate that the use of red blood cells (RBCs) as oxygen carriers and associated secondary hemolysis may in fact cause renal injury, offsetting potential advantages. During 48-hour normothermic perfusion of seven human deceased donor kidneys, we observed progressive hemolysis, leading to iron accumulation in perfusate, tissue, and urine. Untargeted lipidomic analysis revealed profound increases in oxidized phospholipid species in perfused kidneys, pointing towards iron-dependent cell death known as ferroptosis. Next, in twelve additional human kidney perfusions, we demonstrate that either dialysis-based free hemoglobin removal or cell-free perfusion attenuates hemolysis-driven iron accumulation, phospholipid peroxidation, and acute kidney injury. Our findings highlight the pathological role of hemolysis and iron on the kidney, urging restraint in the clinical application of RBC-based kidney perfusion.

## INTRODUCTION

Machine perfusion technologies have emerged as an important new modality in the field of organ preservation and transplantation. To address the growing disparity between the available donor organs and the number of patients on the transplant waiting list (1), significant efforts have been devoted to developing advanced perfusion platforms. These platforms aim to create a window of opportunity to facilitate the assessment, reconditioning, repair and regeneration of deceased donor organs (2).

While normothermic machine perfusion (NMP) techniques for liver transplantation have become routine clinical practice (3, 4), successful translation into the field of kidney transplantation, which constitutes the majority of transplant organs, remains elusive. This raises the question how ex vivo NMP of deceased donor livers has been extended up to 12 days (5), with successful transplantation after 3 days of perfusion (6), whereas studies on normothermic kidney perfusion typically range from 1 to 6 hours, and even then, the first randomized controlled trial was discontinued in the absence of a discernible positive effect (7). This is concerning given the increasing clinical application of normothermic kidney perfusion by organ procurement organizations and highlights the need to critically evaluate whether all organs are equally suited for each preservation method.

In this study, we aimed to elucidate what happens to the kidney graft during normothermic machine perfusion, encompassing a detailed analysis of the multi-day perfusion of 19 deceased donor kidneys. Seven deceased donor kidneys were perfused up to 48-hours. We observed that hemolysis provokes iron accumulation in perfusate and tissue. Through untargeted lipidomic profiling we could confirm the accumulation of oxidized phospholipid species (oxPL), pointing towards iron-dependent cell death known as ferroptosis. Finally, we show that either dialysis-based free hemoglobin removal or cell-free perfusion, an approach we recently demonstrated to support four-day human kidney preservation (8), attenuates hemolysis-driven iron accumulation, phospholipid peroxidation, and acute kidney injury.

## RESULTS

### Study design and donor characteristics

In total, 19 human donor kidneys deemed unsuitable for transplantation were perfused. During the first phase, seven kidneys were perfused up to 48-hours at normothermia with a RBC-based perfusate. In the second phase, an additional twelve human kidneys were included for either dialysis-based free hemoglobin removal during RBC-based perfusion or cell-free perfusion. Donor and RBC characteristics are reported in **Table S1**.

### Kidney perfusion dynamics

A clinical grade perfusion platform was adapted to support prolonged (≥24-hour) preservation (**Figure 1A**). During the first phase, seven kidneys were perfused up to 48 hours. The perfusate was composed of two units of washed red blood cells (RBCs), human serum albumin and DMEM F12 as main components (for full composition see **Table S2**). Of the seven kidneys, five maintained a stable renal blood flow (RBF) throughout the 48-hour perfusion period (**Figure 1B**), whereas Kidney5 and Kidney6 had to be stopped earlier due to graft failure (defined as RBF ≤50 mL/min/100g). After 48 hours, all perfusions were stopped due to progressive disturbances in perfusion dynamics (**Figure S1**). Blinded histologic scoring of periodic acid-Schiff (PAS) staining was performed by a renal pathologist (**Table S3**), demonstrating an increase in tubular vacuolization and decrease in tubular dilation beyond 24 hours of RBC-based perfusion. Cold preservation method (SCS or HMP) prior to warm perfusion had no discernible impact on perfusion dynamics (**Figure S2**).

**Figure 1.**
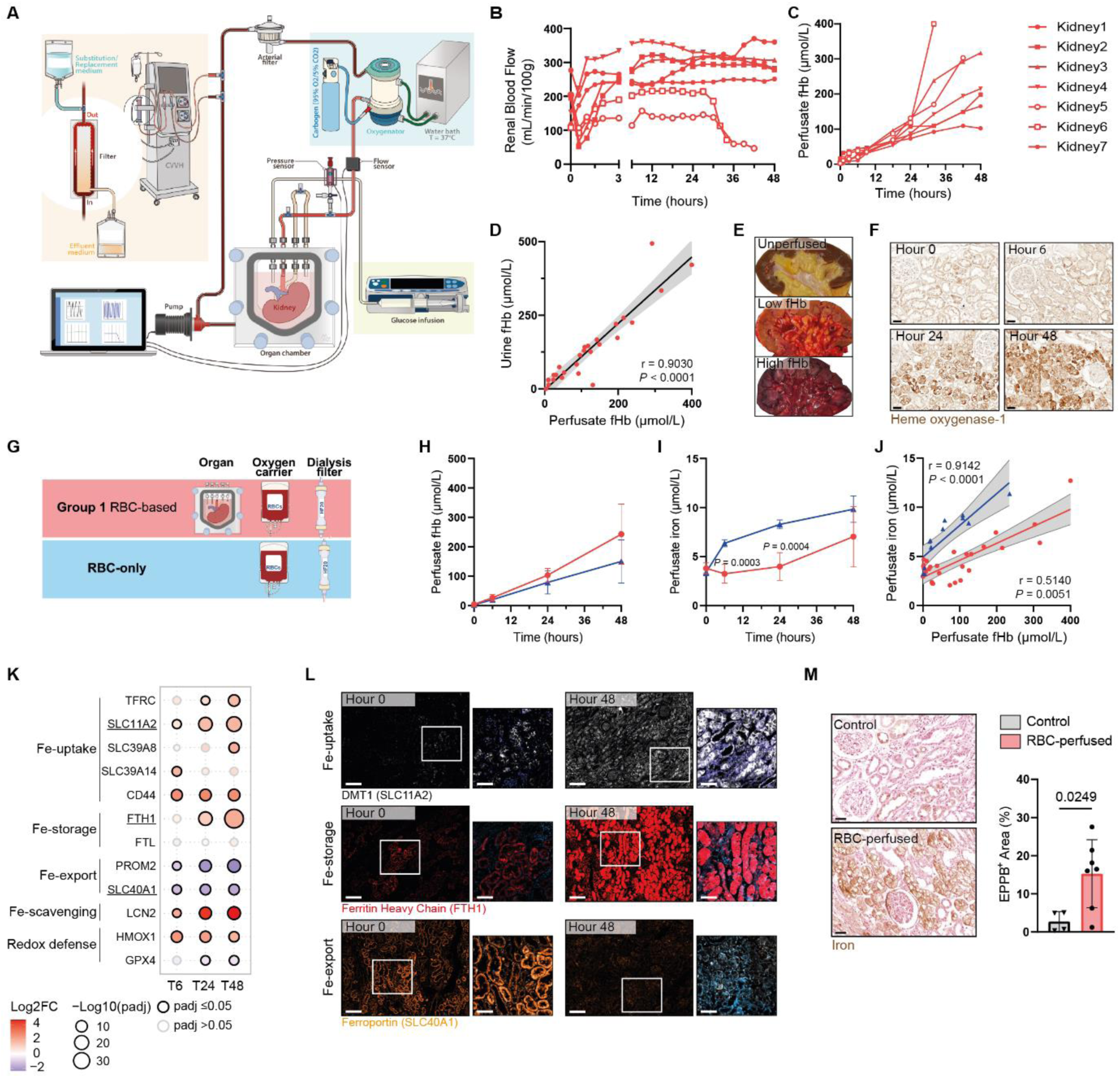
Hemolysis induces iron accumulation during RBC-based human kidney perfusion. **A**, Schematic overview of the perfusion platform. The renal artery was perfused by a pressure-controlled impeller pump set at 75 mmHg with a RBC-based perfusate after it passed through an oxygenator at normothermia. Urine was recirculated. Continuous hemofiltration allowed removal of small molecular weight waste products and substitution with fresh perfusate. **B**, Renal blood flow. **C**, Perfusate free hemoglobin (fHb) accumulated during perfusion. **D**, Association between perfusate fHb and urine fHb (Spearman’s rank correlation coefficient). **E**, Macroscopic appearance of a control kidney, Kidney1 with low fHb, and kidney6 which failed after 34-hours with significant fHb accumulation. **F**, Immunohistochemistry for heme oxygenase-1 (HO1). Representative images of biopsies taken during perfusion. Kidney3 is shown. Scale bar, 50 μm. **G,** Schematic overview illustrating the different experimental groups. Comparison between RBC-based perfusion with (n=7) or without (n=3) a deceased donor kidney in the platform. **H**, Perfusate fHb. **I**, Perfusate iron. Two-way ANOVA. **J**, Association between perfusate fHb and perfusate iron for both groups (Spearman’s rank correlation coefficient). **K,** Gene expression dynamics of key iron uptake, iron storage, iron export, iron scavenging, and redox defence genes during RBC-based human kidney perfusion. Log2 foldchange (Timepoint vs T0) is indicated by colour. Significance (padj ≤0.05) by strong border and size. **L**, Immunofluorescence for DMT1 (Fe-uptake), FHT1 (Fe-storage), and Ferroportin (Fe-export). Representative images of biopsies taken before and after perfusion. Kidney3 is shown. Scale bar, 200 and 50 μm, respectively. **M**, Enhanced Perl’s Prussian blue (EPPB) staining for iron deposition in tissue. Representative images of control (n=4) and RBC-perfused human kidneys (n=7) and quantification (unpaired t-test). Kidney3 is shown. Scale bar, 50 μm.

### Hemolysis drives progressive iron accumulation

Over the course of perfusion we observed the occurrence of hemolysis with buildup of free hemoglobin (fHb) within the perfusate (**Figure 1C**) and associated hemoglobinuria (**Figure 1D**), corroborating earlier preliminary data (9). Distinct macroscopic differences were discernible between a control kidney, Kidney1 which exhibited minimal fHb buildup, and Kidney6 which failed after 34 hours with significant fHb accumulation (**Figure 1E**). Notably, both renal grafts that experienced failure during perfusion were exposed to the highest levels of fHb accumulation.

Heme oxygenase-1 (HO-1), the endogenous defence mechanism against hemolysis-induced kidney injury (10), was upregulated between 6 and 24 hours of perfusion (**Figure 1F**). This enzyme degrades heme into labile iron, carbon monoxide and biliverdin (10), mitigating the deleterious effects of fHb accumulation.

Hemolysis is an inherent consequence of mechanical perfusion of (stored) RBCs (10, 11). Preliminary experiments, in which RBCs were perfused throughout the platform in the absence of a donor kidney, confirmed that the occurrence of hemolysis during prolonged perfusion (≥6 hours) was primarily attributable to the normothermic perfusion system itself, rather than the donor graft (**Figure 1G-J** and **Figure S3**), corroborating previous observations reported in the context of liver perfusion (12, 13). Importantly, comparison of perfusate iron levels during RBC-only perfusion indicates the use of RBCs and subsequent hemolysis as source of circulating iron, with significant iron uptake by the perfused kidney as early as 6 hours into perfusion (**Figure 1I-J**).

Through bulk RNA sequencing on biopsies taken during RBC-based kidney perfusion we assessed gene expression dynamics of key iron uptake, iron storage, iron export, iron scavenging, and redox defence genes (**Figure 1K**). The upregulation of iron uptake and storage proteins, alongside the downregulation of iron export proteins, strongly supports the observed uptake of circulating iron from the perfusate. This was further confirmed at the protein level for divalent metal transporter-1 (DMT1, Fe-uptake), ferritin heavy chain-1 (FTH1, Fe-Storage), and ferroportin (FPN, Fe-export) (**Figure 1L**). Finally, enhanced Perl’s Prussian blue (EPPB) staining indeed demonstrated the accumulation of tissue iron within RBC-perfused kidneys (**Figure 1M**).

We conclude that the use of RBCs, and subsequent hemolysis, during machine perfusion does not only result in renal exposure to high levels of fHb, but also drives cellular accumulation of hemolysis-derived iron.

### RBC-based perfusion triggers ferroptosis

Dysregulated cellular iron homeostasis is known to induce lipid peroxidation, the main characteristic of an iron-dependent type of cell death called ferroptosis (11, 14). In an untargeted approach we sought to characterize changes in the cellular lipidome following RBC-based perfusion, using liquid chromatography tandem mass spectrometry (LC-MS/MS) with subsequent lipid identification through MS-DIAL (15). Overall, 662 unique lipid species were identified, belonging to 11 lipid classes (**Figure 2B** and **Table S4**). To depict differences between control kidneys and those subjected to RBC-based perfusion, we projected relative lipid abundance of the 662 lipid species across these 11 lipid (sub-)classes on a heatmap (**Figure 2C-D**). Our analysis revealed that 50 lipid species exhibited significant increases in perfused kidneys (Log2FC ≥1.5 and a *P* value <0.05; **Table S5**), the majority of which were oxidized phospholipids (oxPL) (**Figure 2E**), indicating significant oxidative modifications during kidney RBC-based perfusion. Normalization to their non-oxidized PL (non-oxPL) pool confirmed significant increases in oxPL/non-oxPL abundance (**Figure 2F-H**). PL peroxidation results in the formation of lipid peroxides, which can further decompose into toxic secondary products, such as malondialdehyde (MDA) and 4-hydroxy-nonenal (4HNE), commonly used as biomarkers for ferroptosis (16). Notably, levels of MDA, assessed as thio-barbituric acid reactive substances (TBARS) in perfusate, and 4HNE in tissue increased significantly over the 48-hour RBC-based perfusion period (**Figure S4**).

**Figure 2.**
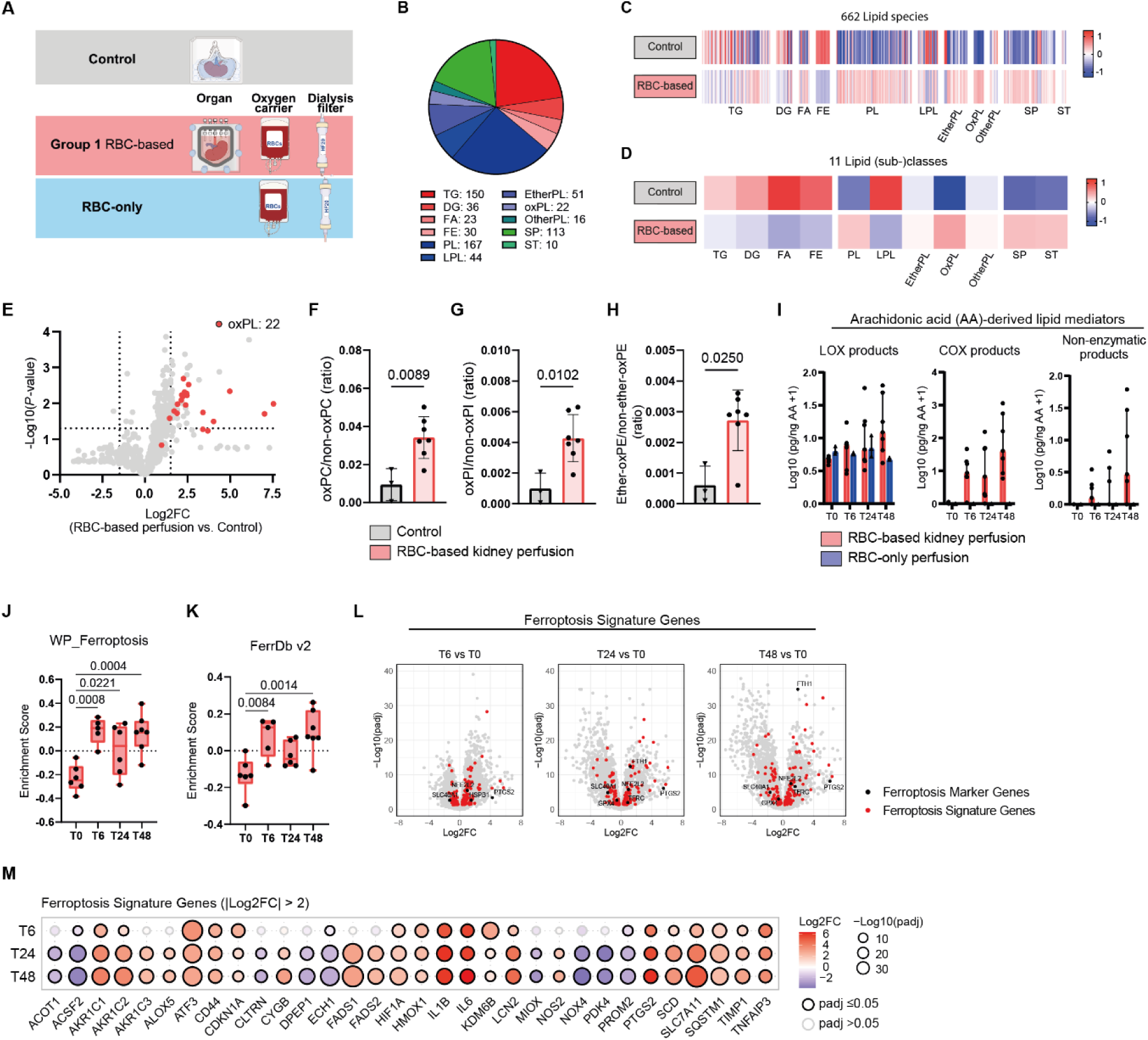
RBC-based human kidney perfusion induces ferroptosis. **A**, Schematic overview illustrating the different experimental groups. **B**, Representation of 662 lipid species detected through untargeted LC-MS/MS lipidomics grouped by lipid (sub-)class (also see **Table S4**). **C-D**, Heatmaps of the relative lipid abundance of the 662 lipids as individual species (**B**) and as 11 lipid (sub-)classes (**C**) for control and RBC-perfused human kidneys (n=3 and n=7, respectively). Z-score represents total area normalized abundance. Abbreviations: TG, Triradylglycerols; DG, Diradylglycerols; FA, Fatty acids; FE, Fatty esters; PL, Phospholipids; LPL, Lyso-phospholipids; EtherPL, Ether-linked phospholipids; oxPL, Oxidized phospholipids; OtherPL, Other phospholipids (BMP (6), CL (9), and MLCL (1)); SP, Sphingolipids; ST, Sterol lipids. **E**, Volcano plot representing the 662 lipid species Log2FC and P-value in RBC-perfused versus control kidneys (unpaired t-test with Welsch correction). Red represents the oxPL species. A list of the lipid species that exhibited significant increases in perfused kidneys can be found in **Table S5** (Log2FC ≥1.5 and P-value <0.05). **F**, Abundance of oxidized PCs/non-oxidized PCs (unpaired t-test). **G**, Abundance of oxidized phosphatidylinositols (PIs)/ non-oxidized PIs (unpaired t-test). **H**, Abundance of ether-linked oxidized phosphatidylethanolamine (PE)/ non-oxidized ether-linked PEs (unpaired t-test). Data are represented as mean with SD. **I**, Levels of arachidonic acid (AA)-derived lipid mediator release into the perfusate of RBC-based kidney perfusion (n=7) and RBC-only perfusion (n=3) at various timepoints. Data are represented as median with IQR. **J**, Box plot of gene set variation analysis enrichment score of WP_Ferroptosis (n=64 genes) (One-way ANOVA). **K**, Box plot of gene set variation analysis enrichment score of FerrDb v2 driver, suppressor and marker genes (n=313 genes) (One-way ANOVA). Data are represented as median with minimum to maximum (**J-K**). **L**, Volcano plots representing significantly up- and downregulated genes (Log2FC and −Log10(padj) at different timepoints during RBC-based kidney perfusion. Black highlights FerrDb v2 marker genes (n=9). Red highlights validated FerrDb v2 driver or suppressor genes with a protein product (n=312). **M**, Dotplot illustrating dynamics in FerrDb v2 driver, suppressor, and marker genes during RBC-based kidney perfusion relative to baseline (|Log2FC|≥2 and padj ≤ 0.05 for at least one of the timepoints).

As hemolysis occurs already in the beginning of perfusion we next assessed dynamics in phospholipid peroxidation during perfusion, for which oxilipidomic analysis (LC-MS/MS) was performed on perfusate taken before connection of the kidney (T0), and after 6, 24, and 48 hours. This enables differentiation between enzymatic phospholipid peroxidation – mediated by lipoxygenases (LOXs) and cyclooxygenases (COXs) – and non-enzymatic lipid peroxidation, which primarily occurs via Fenton reaction (17). Comparison of RBC-only with RBC-based kidney perfusion revealed presence of LOX-derived products across all timepoints in both experimental conditions (**Figure 2I**). Conversely, COX-derived products and non-enzymatic oxPLs-derived products were absent during RBC-only perfusion, and progressively increased over the course of RBC-based kidney perfusion (**Figure 2I**).

Significant enrichment of ferroptosis gene sets was observed already 6 hours into RBC-based perfusion (**Figure 2J-K**). Next, we compared differentially expressed genes (DEGs) across timepoints, specifically focusing on ferroptosis marker, driver and suppressor genes (as defined by FerrDb v2 (18)) (**Figure 2L**), for which the genes with the most significant dynamics (|Log2FC|≥2 and padj≤0.05) for at least one of the timepoints are visualized in **Figure 2M**.

We conclude that the progressive accumulation of oxidized phospholipids, coupled with the enrichment of ferroptosis-related gene sets, points towards ferroptotic cell death as a result of hemolysis-driven iron accumulation during RBC-based kidney perfusion.

### Interventions to mitigate hemolysis-driven renal injury

Having established that progressive hemolysis during machine perfusion leads to renal iron accumulation and the buildup of oxPL species, we questioned whether these effects could be mitigated. In the second phase of this study, we perfused an additional twelve human kidneys, assigning them to two intervention groups: fHb removal during RBC-based perfusion (Group2) and cell-free perfusion (Group3).

### Free hemoglobin removal reduces phospholipid peroxidation

With the first intervention (Group2), we aimed to remove fHb from the perfusate by replacing the HF20 dialysis filter with the Oxiris filter, a novel blood purification filter for continuous renal replacement therapy that removes fHb (**Figure 3A**). Four kidneys were perfused with the Oxiris filter up to 48-hours, showing comparable perfusion dynamics and histology to the RBC-perfused kidneys (**Figure S5** and **Table S3**). Dialysis-based fHb removal significantly decreased perfusate fHb levels (**Figure 3B-C**), with substantial clearance into the effluent (**Figure 3D**). This was accompanied by a trend towards reduced tissue iron accumulation when compared to RBC-perfused kidneys (**Figure S6**).

**Figure 3.**
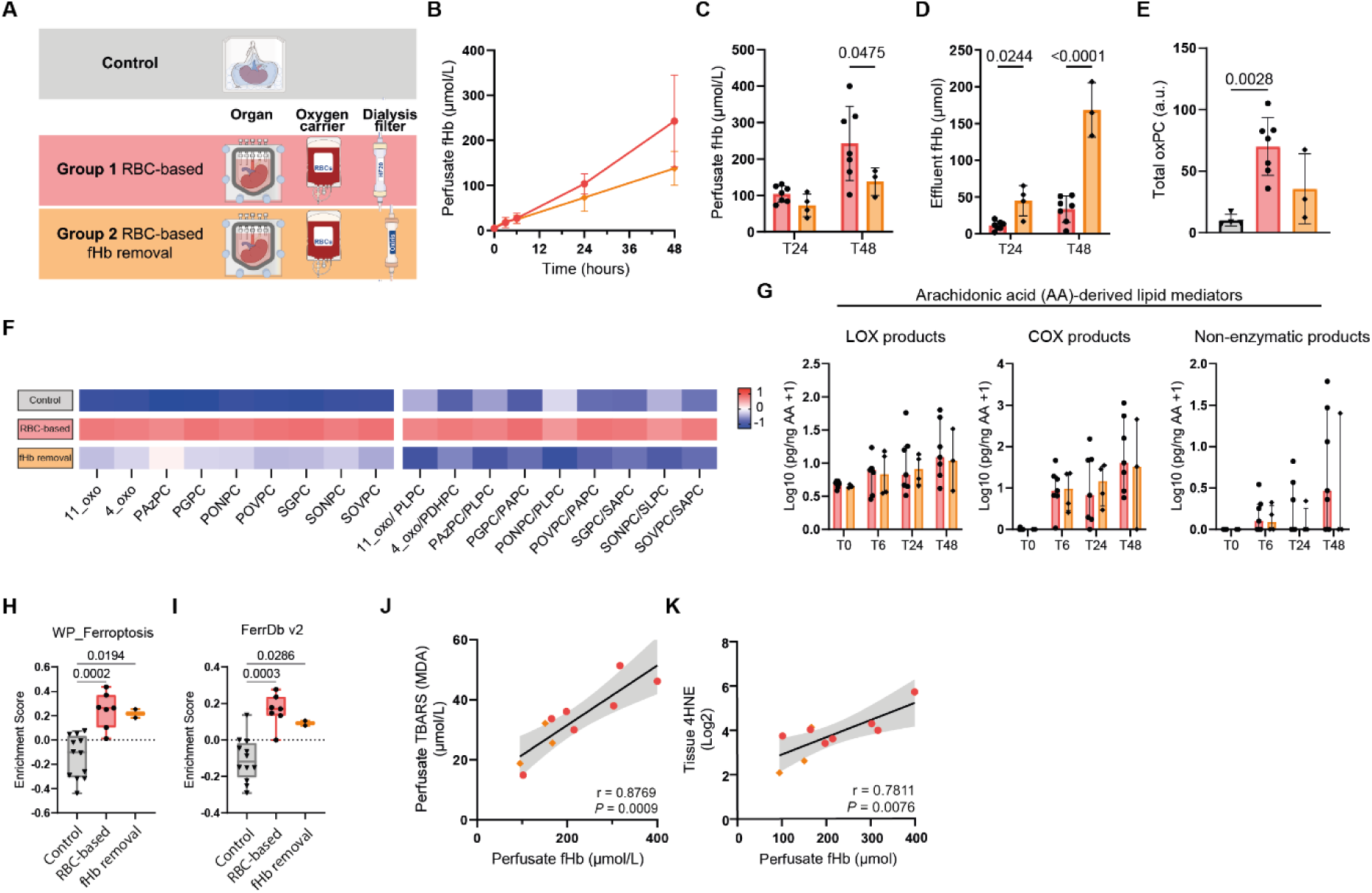
Dialysis-based removal of free hemoglobin reduces hemolysis-driven phospholipid peroxidation. **A**, Schematic overview illustrating the different experimental groups. Comparison between dialysis-filter based free hemoglobin (fHb) removal during RBC-based perfusion (Group2, n=4) with RBC-based perfusion (Group1, n=7) of deceased human donor kidneys. **B-D**, Perfusate fHb during perfusion (**B**), and perfusate (**C**) and effluent (**D**) fHb after 24 and 48 hours of perfusion (Two-way ANOVA). **E**, Total oxidized phosphatidylcholines (oxPC) abundance as detected through targeted oxPC LC-MS/MS (One-way ANOVA). **F**, Heatmap of the relative lipid abundance for the different oxPC as detected through targeted oxPC analysis. See **Table S6** for identification of the different PC and oxPCs and their abbreviations. **G**, Levels of arachidonic acid (AA)-derived lipid mediator release into the perfusate during perfusion. Data are represented as median with IQR. **H**, Box plot of gene set variation analysis enrichment score of WP_Ferroptosis (n=64 genes) (One-way ANOVA). **I**, Box plot of gene set variation analysis enrichment score of validated FerrDb v2 driver, suppressor and marker genes (n=313 genes) (One-way ANOVA). Data are represented as median with minimum to maximum (**H-I**). **J**, Association between perfusate fHb and perfusate TBARS (MDA) after 48 hour perfusion (Pearson’s correlation coefficient). **K**, Association between perfusate fHb and tissue 4-hydroxynonenal (4HNE) after 48 hour perfusion (Pearson’s correlation coefficient).

Through targeted LC-MS/MS oxidized phosphatidylcholines (oxPC) analysis on tissue at the end of preservation, we observed that the oxidative modifications during the RBC-based perfusions were reduced through fHb removal (**Figure 3E-F**). However, oxilipidomic profiling revealed continued release of LOX-, COX-, and non-enzymatic oxidation products into the perfusate (**Figure 3G**), indicating persistent ferroptotic activity. This was further supported by unaffected enrichment of the ferroptosis-related gene signatures following fHb removal (**Figure 3H-I** and **Figure S6**).

Hence, while fHb removal attenuates phospholipid peroxidation, it does not completely prevent it, likely due to the continued presence of fHb in the perfusate, albeit at reduced levels. Notably, despite fHb removal during RBC-based perfusion, MDA (measured as TBARS) in perfusate and 4HNE in tissue still demonstrated a strong correlation with perfusate fHb and iron levels, highlighting the persistent pro-oxidative effects of residual hemolysis products (**Figure 3J-K** and **Figure S6**).

We conclude that, even after fHb removal, residual hemolysis continues to contribute to lipid peroxidation, albeit at lower levels when compared to RBC-based perfusion.

### Cell-free perfusion evades hemolysis-driven iron accumulation and ferroptosis

The most definitive approach to avoid hemolysis, which would be the removal of RBCs from the perfusate altogether, was tested in the second intervention group (Group3) (**Figure 4A**). Lowering the perfusion temperature from 37°C to 25°C allows for cell-free perfusion as it reduces the metabolic rate, and therewith oxygen requirements. As we previously demonstrated, this cell-free perfusion approach enables four-day ex vivo metabolic and functional preservation of human kidneys (8). Eight kidneys were preserved with a cell-free perfusate, which confirmed improved metabolic preservation during 48-hour perfusion when compared to RBC-based perfusion (**Figure S7**).

**Figure 4.**
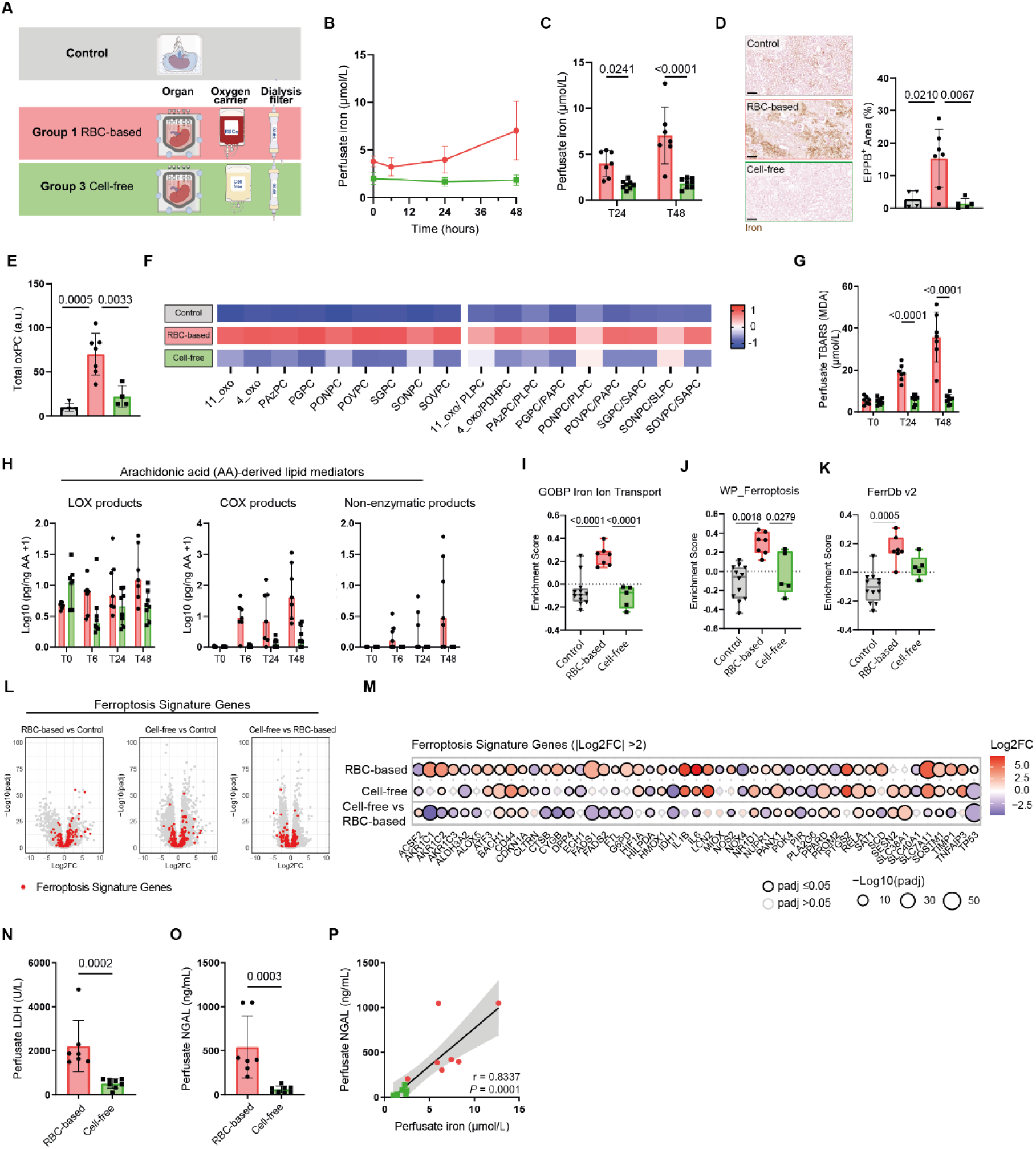
Cell free perfusion negates hemolysis-driven iron accumulation and phospholipid peroxidation. **A**, Schematic overview illustrating the different experimental groups. Comparison between cell-free perfusion (Group3, n=8) with RBC-based perfusion (Group1, n=7) of deceased human donor kidneys. **B-C**, Perfusate iron during perfusion (**B**) and perfusate iron after 24 and 48 hours of perfusion (**C**) (Two-way ANOVA). **D**, Enhanced Perl’s Prussian blue (EPPB) staining for iron deposition in tissue. Representative images and quantification (One-way ANOVA). Scale bar, 200 μm. **E**, Total oxidized phosphatidylcholines (oxPC) abundance as detected through targeted oxPC LC-MS/MS (One-way ANOVA). **F**, Heatmap of the relative lipid abundance for the different oxPC as detected through targeted oxPC analysis. See **Table S6** for identification of the different PC and oxPCs and their abbreviations. **G**, Perfusate TBARS (MDA) after 0, 24, and 48 hours of perfusion (Two-way ANOVA). **H**, Levels of arachidonic acid (AA)-derived lipid mediator release into the perfusate during perfusion. Data are represented as median with IQR. **I**, Box plot of gene set variation analysis enrichment score of GOBP_Iron_Ion_Transport (n=58 genes) (One-way ANOVA). **J**, Box plot of gene set variation analysis enrichment score of WP_Ferroptosis (n=64 genes) (One-way ANOVA). **K**, Box plot of gene set variation analysis enrichment score of validated FerrDb v2 driver, suppressor and marker genes (n=313 genes) (One-way ANOVA). Data are represented as median with minimum to maximum (**I-K**). **L**, Volcano plots representing significantly up- and downregulated genes (Log2FC and −Log10(padj) after RBC-based or cell-free kidney perfusion, compared to control kidneys, and cell-free perfusion compared to RBC-based perfusion. Red highlights validated FerrDb v2 driver, suppressor, or marker genes with a protein product (n=313). **M**, Dotplot illustrating dynamics in FerrDb v2 driver, suppressor, and marker genes after RBC-based or cell-free perfusion, compared to control kidneys, and cell-free perfusion compared to RBC-based perfusion (|Log2FC|≥2 and padj ≤ 0.05 for at least one of the comparisons). **N**, Perfusate LDH after 48-hour perfusion (Mann-Whitney test). **O**, Perfusate NGAL after 48-hour perfusion (Mann-Whitney test). **P**, Association between perfusate NGAL and iron (Spearman’s rank correlation coefficient).

Perfusate iron levels did not increase during 48-hour cell-free perfusion, in contrast to RBC-based perfusion. (**Figure 4B-C**). Similarly, tissue iron accumulation, as observed in RBC-perfused kidneys, was absent during cell-free perfusion (**Figure 4D**). This confirms that the use of RBCs, and subsequent hemolysis, drives iron accumulation during RBC-based kidney perfusion. Significantly lower perfusate bilirubin levels following cell-free perfusion further point towards a role of HO-1 mediated breakdown of fHb into labile iron (**Figure S7**).

Targeted oxPC analysis revealed that oxidative modifications observed after RBC-based perfusions were attenuated by cell-free perfusion (**Figure 4E-F**). In line with this, perfusate TBARS levels did not increase during cell-free perfusion (**Figure 4G**). Oxilipidomic profiling in the perfusate showed the presence of LOX- and COX-derived oxidation products during cell-free perfusion, with COX-derived products being less abundant as compared to RBC-based perfusion. Importantly, non-enzymatic oxidation products were undetectable in the perfusate during 48-hour cell-free kidney perfusion (**Figure 4H**), supporting the hypothesis that free hemoglobin-derived iron plays a critical role in Fenton-driven lipid peroxidation during RBC-based kidney perfusion.

Although the cell-free perfusion group included both DCD and DBD donors, whereas the RBC-perfused group only contained DCD donors, donor background had a negligible impact on perfusion outcomes (**Figure S8**).

The distinct differences observed in tissue iron accumulation and lipid peroxidation between RBC-based and cell-free perfusion were corroborated at the transcriptomic level, where cell-free perfusion showed markedly reduced enrichment of iron ion transport and ferroptosis-related gene sets (**Figure 4I-K**). Next, differential gene expression analysis further highlighted differences in ferroptosis marker, driver and suppressor genes between RBC-based and cell-free perfusion (as defined by FerrDb v2 (18)) (**Figure 4L**), for which the genes with the most significant changes (|Log2FC|≥2 and padj≤0.05) for at least one of the timepoints are visualized in **Figure 4M**.

The absence of iron accumulation and subsequent lipid peroxidation during cell-free perfusion underscore that RBCs, and subsequent hemolysis, drive iron accumulation and ferroptosis during RBC-based kidney perfusion. Finally, we assessed biomarkers for acute kidney injury (AKI) during perfusion, observing significantly higher levels of perfusate lactate dehydrogenase (LDH) and neutrophil gelatinase associated lipocalin (NGAL) in RBC perfused kidneys (**Figure 4N-O**). This is striking given that two donor kidneys in the cell-free perfusion group had significant dysfunction at the time of donation (as indicated by a final serum creatinine >200 µmol/L) (**Figure S9**).

As NGAL is an iron-sequestering siderophore, these findings raise the possibility that NGAL is produced by the kidney during RBC-based perfusion in response to hemolysis to protect the kidney from iron-induced injury. Indeed, we did find a strong correlation between perfusate iron and NGAL (**Figure 4P**), which is in line with experimental studies that have suggested that NGAL may play a protective role during AKI (19). Whereas NGAL is a kidney-specific marker, LDH can also originate from hemolysis. However, supporting experiments confirmed that the observed increases are predominantly kidney-derived (**Figure S2**), confirming that hemolysis during RBC based kidney perfusion exacerbates renal injury.

## DISCUSSION

In this study, we observed that the use of RBCs and associated hemolysis during machine perfusion of diseased donor kidneys drives progressive iron accumulation and phospholipid peroxidation. The observed accumulation of fHb, iron, oxidized phospholipids, and their byproducts highlights how hemolysis within preservation platforms leads to ferroptosis. Finally, we demonstrate that dialysis-based fHb removal reduces, and cell-free perfusion circumvents, hemolysis-driven iron accumulation, phospholipid peroxidation, and acute kidney injury.

Ferroptosis is an iron-dependent form of cell death that is distinguished from other types of cell death by the occurrence of excessive phospholipid peroxidation (11, 20), and has been found to play a central role in renal ischemia reperfusion injury, maladaptive repair, and the development of kidney fibrosis (14, 21–23). Unlike the liver, within which resident macrophages effectively and on-demand manage the removal of damaged RBCs and fHb, and therewith resilience against hemolysis (12, 24, 25), the kidney is particularly vulnerable to elevated levels of fHb and the resultant hemoglobinuria (10, 11, 26). Preclinical studies have emphasized the importance of attenuating iron and lipid peroxide accumulation in the kidney, as this propagates ferroptotic cell death and culminates in AKI (22, 27). Various clinical investigations have demonstrated that hemolysis during cardiopulmonary bypass surgeries (bypass time 1-3 hours) is associated with AKI, need for renal replacement therapy, and even mortality (28, 29).

Although our analysis primarily focused on prolonged preservation (≥6 hours), iron uptake from the perfusate, a ferroptosis gene signature, and release of phospholipid peroxidation products was already observed after 6 hours of RBC-based perfusion. This is concerning given an increasing number of clinical trials looking into normothermic kidney perfusion, and clinical implementation of kidney NMP by commercial organ procurement organizations.

Machine perfusion of deceased donor organs is a rapidly developing field that holds the potential to change transplantation medicine. Despite the prevailing assumption that RBC-based NMP confers protection to deceased donor kidneys (2, 30), our findings challenge this notion by demonstrating that the use of RBCs as oxygen carriers, and associated hemolysis, may actually offset the anticipated benefits of normothermic perfusion. Our study underscores the pathological role of hemolysis and iron in exacerbating AKI. Consequently, we urge caution in the clinical implementation of RBC-based perfusion for deceased donor kidneys. Our findings challenge the current kidney preservation dogma, and necessitate reconsideration of the framework needed to establish a protective environment to facilitate ex situ reconditioning, repair, and regeneration of deceased donor kidneys.

## METHODS

### Ethics

This research complies with all relevant ethical regulations. Leiden University Medical Center (LUMC) received authorization from the Dutch government (BWBR0008974, 2555663-CZ/IZ/2562427) for kidney transplantation and associated research. Prior to organ retrieval for all human kidneys, donor research consent was obtained by Eurotransplant, the centralized donation organization in The Netherlands (BWBR008066, Art13). This consent was acquired by an independent organ donation coordinator unaffiliated to the research team.

#### Study design

Nineteen kidneys from deceased donors were subjected to multi-day perfusion and allocated into three experimental groups: seven kidneys were perfused with a red blood cell (RBC)-based perfusate at normothermia (37°C) (“RBC-based”; Group1), four kidneys were perfused with a similar RBC-based perfusate at normothermia but incorporated a dialysis filter that allowed removal of free hemoglobin (“fHb removal”; Group2), and eight kidneys were perfused with a cell-free perfusate (i.e. no RBCs) at subnormothermia (25°C) (“cell-free”; Group3). Tissue biopsies obtained at the end of perfusion were compared to biopsies taken from control deceased donor kidneys and assessed through both untargeted lipidomics and targeted oxidized phosphatidylcholines (oxPC) analysis. Additionally, three perfusions were performed using a RBC-based perfusate at normothermia in the absence of a donor kidney (“RBC-only”) to assess the interaction between the perfusion platform, stored RBCs, and deceased donor kidneys.

#### Perfusion platform and protocol

*Adapted perfusion platform*. A clinical-grade perfusion platform (CirQlife, XVIVO) was modified to support prolonged (≥24-hour) preservation. This adapted platform consists of a closed-loop circuit connected to a custom-designed 3D printed organ chamber (Oceanz) that also serves as venous reservoir. A pressure-controlled impeller pump (CirQlife, XVIVO) perfused the renal artery through silicone tubing (LS17 or LS25, Masterflex Metrohm) at a mean arterial pressure (MAP) of 75 mmHg (60 BPM, 20% amplitude). Pressure in the renal artery was measured in-line with a single-use pressure sensor (Edwards Lifesciences). Flow was monitored using an ultrasonic clamp-on flow probe (Transonic Systems Europe B.V.), and temperature was monitored using a temperature sensor (Heraeus Nexensos PT100 Surface Sensor [W-SZK], Heraeus Nexensos GmbH), which was positioned underneath the organ. Arterial pressure, perfusate flow and temperature were continuously recorded. The perfusate was oxygenated with a carbogen mixture of 95% O2 and 5% CO2 through a long-term membrane oxygenator (Lilliput 2 ECMO, LivaNova) that was connected to a water bath set at set at 37 °C. Urine was recirculated.

A Prismaflex system (Baxter) was connected in parallel to remove metabolic waste products and supply fresh nutrients whilst maintaining electrolyte homeostasis. A paediatric filter (Prismaflex HF20 set, Baxter) allowed the exchange of small molecular weight molecules. Blood flow over the filter was set at 20 mL/ min. Fluid and small molecular weight molecules were removed at 80 mL/ hr over the filter. Fresh substitution solution (see **Table S2** for composition) was substituted post-filter at the same rate (filtration fraction of 10%). A separate infusion pump infused additional glucose (5%, WE0087G, Baxter) at a rate of 0-7 mL/ hr to maintain glucose above 4 mmol/ L.

*Human kidneys.* After in situ flushing of the with cold University of Wisconsin (Belzer UW, Bridge to Life Ltd.) preservation solution, the kidneys were retrieved and transported to the LUMC by static cold storage (SCS) or hypothermic machine perfusion (HMP). Upon arrival at the laboratory, the renal artery and ureter were cannulated using Luer lock connectors (Cole Parmer) whilst the perfusion platform was setup and primed in parallel. Shortly before the start of culture kidneys were flushed with DMEM F12 supplemented with human serum albumin (HSA) (40g/L).

*Perfusate.* A detailed description of the perfusate composition is provided in **Table S2**. Two units of stored clinical-grade leukocyte-depleted RBCs (Sanquin) were used per perfusion. Prior to their addition, stored RBCs were washed three times with DMEM F12 through centrifugation at 1000g for 10 minutes (21°C, acc9, dec4) to remove RBC storage solution and hemolytic RBCs (9, 31).

In parallel, the perfusion platform was primed with the other perfusate components, after which the washed RBCs were also added. Depending on perfusate acidity, pH was adjusted with sodium bicarbonate (7.5%, Gibco) until within range (Target 7.25-7.40). Perfusion was started with a volume of approximately 1L.

*Intervention group : free hemoglobin (fHb) removal*. The paediatric HF20 filter was exchanged for an Oxiris dialysis filter (Baxter), allowing removal of free hemoglobin into the effluent (dialysis waste). fHb levels were measured in both perfusate and effluent over the course of perfusion. The substitution solution remained unchanged. Blood flow over the filter was set at 20 mL/ min. Fluid was removed at 100 mL/ hr over the filter, with fresh substitution solution being supplemented post-filter at the same rate (filtration fraction of 13%). Of the four human kidneys perfused in this group, one was stopped after 24 hours of perfusion due to logistical reasons.

*Intervention group : cell-free perfusion*. Kidneys were perfused using our previously described cell-free perfusion approach, which enabled four-day ex vivo metabolic and functional preservation of human kidneys at subnormothermia (25°C) (8). Lowering perfusion temperature from 37°C to 25°C allows for cell-free perfusion as it reduces the metabolic rate, and therewith oxygen requirements. Hence, hemolysis cannot occur and fHb is absent.

*Control group : RBC-only perfusions*. For the three perfusions performed in the absence of a kidney graft, we followed the same procedure as for RBC-perfused kidneys, except for the donor kidney being absent. The resistance normally provided by the kidney was simulated by partially clamping the tubing at the place where the renal artery is normally connected, thereby creating a flow rate and vascular resistance similar to those observed during RBC-based kidney perfusions.

### Assessment of kidney preservation and injury

Perfusate was sampled at least three times per day throughout the perfusion period. Arterial, venous and urine samples were measured directly for blood-gas analyses and monitoring of electrolytes and metabolites using an iSTAT1 blood analyzer (Abbott) and CG4^+^ (pH, pO_2_, pCO_2_, HCO ^−^, lactate, and saturation) and Chem8 iSTAT cartridges (sodium, potassium, chloride, glucose, and Hb) (Abott). Oxygen uptake (mL O_2_/ min/ 100gr) was calculated as:

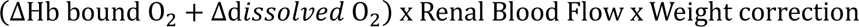

Perfusate samples were centrifuged at 1000g for 7 minutes (4°C, acc9, dec4), whereafter the supernatant was aliquoted and frozen at −80 °C for later analysis. Effluent samples (i.e. fluid removed through continuous hemofiltration) was sampled after 24 and 48 hours.

The Clinical Chemistry Laboratory within the LUMC measured perfusate, urine, and effluent free hemoglobin (fHb), albumin, total bilirubin (tBil), total iron, LDH, and ferritin according to standard operating procedures, that had previously been stored at −80°C.

Neutrophil gelatinase-associated lipocalin (NGAL) and kidney injury molecule-1 (KIM1) levels in the perfusate were measured using a quantitative sandwich enzyme immunoassay technique with NGAL Quantikine ELISA kit (DLCN20, R&D systems) and KIM-1 DuoSet ELISA kit (DY1750B, R&D systems) according to manufacturer’s instruction in samples that had previously been frozen and stored at −80 °C. A thiobarbituric acid reactive substances (TBARS) assay kit (CaymanChemical, 10009055) was used to measure levels of malondialdehyde (MDA) and related lipid oxidation end products in the perfusate according to manufacturer’s instructions in samples that had previously been frozen and stored at −80 °C.

### Tissue sampling and processing

Wedge biopsies were taken at the end of perfusion and from control deceased donor kidneys and either fixed in 4% formaldehyde (VWRK4187F, VWR) or snap frozen in liquid nitrogen and stored at −80 °C for further analysis. Punch biopsies (6 mm) were taken at four timepoints during perfusion (Hour 0, Hour 6, Hour 24, and Hour 48). At each timepoint biopsies were cut longitudinally into two pieces of which one was fixed in formaldehyde and the other snap frozen in liquid nitrogen. Paraffin-embedded tissue sections were sectioned at 5 µm for histochemical and immunohistochemistry staining.

### Histochemical staining

For Enhanced Prussian Blue staining (EPPB), paraffin-embedded tissue sections were dewaxed and stained with an iron stain kit (ab150674, Abcam). Human spleen was stained as positive control. To enhance iron detection, we adapted the EPPB staining protocol as described previously (32), by adding a blocking step (Protein block, s1699, Dako) for 60 minutes followed with 30 minutes of peroxidase blocking solution (ab64218, Abcam) and a 30 minute incubation with 3,3-diaminobenzidine (DAB) substrate (ab64238, Abcam) on sections that had been previously stained with the iron stain kit. A counterstain with nuclear fast red solution was performed for 2 minutes. Samples were dehydrated and mounted with Entellan (1079610100, Merck). Whole slides were digitized using a 3D Histech Pannoramic MIDI Scanner (Sysmex) and viewed with CaseViewer software. A minimum of 7 Fields of View (3870 x 2050 µm) were selected per sample. Prussian blue positive area was quantified in ImageJ using the H DAB plugin after correction for background staining. Periodic acid-Schiff (PAS) staining was performed on paraffin-embedded tissue sections by the Pathology department using an automatic slide stainer. Slides were digitized and scored by a blinded renal pathologist on acute tubular injury, tubular casts, tubular vacuolization, tubular dilation, and interstitial edema.

### Immunohistochemistry

For immunohistochemistry, paraffin-embedded tissue sections were dewaxed. Antigen retrieval was performed using antigen retrieval buffer (s1699, Dako) in an autoclave, after which a 60 minutes protein block (X0909, Dako) and 15 minutes peroxidase block (ab64218, Abcam) were performed. Primary anti-HO1 (MA1-112, ThermoFisher) and anti-4HNE (MA5-27570, ThermoFisher) and isotype mouse IgG1 (14-4714-82, Invitrogen) were used. All primary antibodies and isotypes were incubated at 4 µg/mL overnight at 4°C in 1%NDS/ 1% NGS/ 1%BSA in PBS. Thereafter, samples were incubated for 1-2 hours at RT with goat anti-Mouse HRP (2 µg/mL, P0447, Dako) followed by incubation with DAB substrate (ab64238, Abcam) for approx. 10 minutes. A counterstain with Mayer’s hematoxylin was performed for 40 seconds. Samples were dehydrated and mounted with Entellan (1079610100, Merck). Whole slides were digitized using a 3D Histech Pannoramic MIDI Scanner (Sysmex) and viewed with CaseViewer software.

### Immunofluorescence

For immunofluorescence, paraffin-embedded tissue sections were dewaxed. Antigen retrieval was performed using antigen retrieval buffer (s1699, Dako) in an autoclave, followed by a 60 minutes protein block (X0909, Dako). Primary anti-DMT1 (Ab55735, Abcam), anti-FTH1 (701934, Invitrogen), and anti-ferroportin (PA5-2293, ThermoFisher) and isotype mouse IgG2a (X0943, DAKO) and rabbit IgG (X0936, DAKO) were used. All primary antibodies and isotypes were incubated at 5 µg/mL overnight at 4°C in PBS. Thereafter, samples were incubated for 1-2 hours at RT with goat anti-Mouse IgG2a A647 (4 µg/mL, A-21241, ThermoFisher) or goat anti-rabbit IgG A568 (4 µg/mL, A-11011, ThermoFisher). A counterstain with Hoechst 33258 (1:10.000 in MQ, H3569, Life Technologies) was performed for 5 minutes. Samples were mounted with Prolong Gold (P36930, Invitrogen). Whole slides were digitized using a Zeiss Axioscan 7 microscope slide scanner (Zeiss) and viewed with Zen3.4.

### Western blotting

Frozen kidney tissues were sectioned on a Cryostar NX70 cryostat (ThermoFisher) at 20 µm, with approx. 30 sections per kidney being collected in Eppendorf tubes, and dissolved in RIPA buffer (89900, ThermoFisher) freshly supplemented with protease and phosphatase inhibitors (78440, ThermoFisher) by pipetting. Protein content was determined through Pierce BCA assay kit (23227, ThermoFisher), and lysates were diluted to a concentration of 2222 µg/ mL. Kidney lysate (27 µL) and Laemmli sample buffer (9 µL; 1610747, BioRAD) were combined, and boiled for 5 minutes at 100°C. Then, 30 µL of lysates per lane (equal to 50 µg protein) and a ladder (1610374, BioRAD) were resolved on 10% Mini-PROTEAN TGX Precast Protein gels 4569034, BioRAD) and transferred onto 0.2 µm PVDF membranes (1704156, BioRAD), using Trans-Blot Turbo Transfer System (BioRAD). For blocking and antibody incubation steps, 5% skim milk (70166, Merck) in PBS was used. All washing steps were performed with 0.05% Tween 20 (P1379, Merck) in PBS. Blots were blocked overnight at 4°C, incubated overnight at 4°C with anti-4HNE (1 µg/ mL, MA5-27570, ThermoFisher) or for 1-2 hours at room temperature (RT) with anti-ACTB (0.4 µg/ mL, MA5-15739, ThermoFisher), and subsequently incubated with anti-Mouse HRP antibodies (0.02 µg/ mL, P0447, DAKO) for 1-2 hours at RT. For substrate visualization, SuperSignal^TM^ West Femto (34095, ThermoFisher) and SuperSignal^TM^ West Pico PLUS (34580, ThermoFisher) were used, respectively. Stripping buffer (21059, ThermoFisher) was used between 4HNE and ACTB. Blots were imaged with a ChemiDoc^TM^ Touch imaging system (BioRAD) and analyzed using Image Lab (v5.2). For densitometry, OD of the protein of interest relative to the housekeeping gene was normalized to the averages obtained in control samples.

### Sample preparation of kidney tissue samples for untargeted lipidomics analysis

Frozen kidneys were homogenized in chilled LC-MS-grade water to obtain a final concentration of 100 mg/mL by means of a Bullet BlenderTM 24 (NY, USA, Next Advance Inc.) and 0.9-2.0 mm diameter stainless steel beads. To 25 µL of kidney tissue homogenate (equal to 2.5 mg of frozen tissue), 600 µL of methyl-tert-butyl ether (MTBE) and 150 µL LC-MS grade methanol were added. Then samples were vortexed and centrifuged at 18000×g for 5 min and supernatants were transferred to new Eppendorf tubes. The extraction was repeated with 300 µL MTBE and 100 µL methanol and after centrifugation, supernatants were combined. 300 µL of LC-MS grade water were added and after centrifugation at 18000×g for 5 min, the upper organic layer were transferred to a glass vial and evaporated under a gentle stream of N_2_. Dried samples were stored at −80 °C until the day of analysis. On the day of analysis, samples were reconstituted in 100 µL 2-propanol, vortexed and sonicated for 5 min. 100 µL of LC-MS grade water were added and samples were vortexed and sonicated for 5 min again. The samples were transferred to inserts and were ready to be analysed by the LC-MS/MS untargeted lipidomics platform. A quality control (QC) was made by pooling equal aliquots of every single sample and was analysed periodically during the analytical run.

### Untargeted lipidomic analysis by LC-MS/MS

10 µL of the lipid extracts were injected in a LC-MS/MS equipment composed of a Shimadzu Nexera X2 (Shimadzu) LC system using as eluents water:acetonitrile 80:20 (mobile phase A) and water:2-propanol:acetonitrile 1:90:9 (mobile phase B), both containing 5 mM ammonium formate and 0.05% formic acid. The applied gradient, at a 300 µL/min flow rate was: 0 min 40% B, 10 min 100% B, 12 min 100% B using a Phenomenex Kinetex C18, 2.7 µm particle, 50 × 2.1 mm (Phenomenex) column and a Phenomenex SecurityGuard Ultra C8, 2.7 µm, 5 × 2.1 mm cartridge (Phenomenex) guard column, both kept at 50 °C. The MS system was a Sciex TripleTOF 6600 (AB Sciex Netherlands B.V.) operated both in positive and negative electrospray (ESI) mode, with the following conditions: ion source gas 1 45 psi, ion source gas 2 50 psi, curtain gas 35 psi, source temperature 350 °C, acquisition range *m/*z 100-1800, ion spray voltage 5500 and −4500 V for ESI+ and ESI-, respectively, declustering potential 80 V (ESI+) and −80 V (ESI-). An information dependent acquisition (IDA) strategy was used to identify lipids. The following parameters were used for MS: collision energy ±10 V, acquisition time 250 ms and for MS/MS these were: collision energy ±45 V, collision energy spread 25 V, ion release delay 30, ion release width 14 and acquisition time 40 ms. The IDA switching criteria were set so that only a maximum number of 20 ions were taken into account these being greater than *m/z* 300 exceeding 200 cps and excluding the former target for 2 s and the isotopes within 1.5 Da.

MS-DIAL (v5.2.240424.3) with the associated standard lipid library belonging to this version (15, 33) was used to align the data and identify the different lipids based on their MS/MS fragmentation pattern. The MS-DIAL parameters were as follows: all peaks with an intensity of at least 500 ions eluting from 0.5 to 10 min of the chromatogram were considered. Alignment within samples was performed with a retention time tolerance of 0.15 min and a MS tolerance of 0.025 Da. The following parameters were set as identification settings: 0.01 and 0.025 Da for both MS1 and MS2 accurate mass tolerance, respectively, and including as adducts [M+H]^+^, [M+Na]^+^, [M+K]^+^, [M+NH_4_]^+^ and [M-H_2_O+H]^+^ in ESI+ and [M-H]^−^, [M+Cl]^−^, [M+HCOO]^−^ and [M-H_2_O-H]^−^ in ESI-. Manual curation for all lipid classes was performed by only including those lipids for which the experimental and reference MS/MS spectra matched.

### Materials for targeted oxidized PCs analysis

All reagents used were of LC-MS grade or higher. Non-oxidized PC standards included 1-palmitoyl-2-linoleoyl-*sn*-glycero-3-phosphocholine (PLPC), 1-palmitoyl-2-arachidonoyl-*sn*-glycero-3-phosphocholine (PAPC), 1-stearoyl-2-linoleoyl-*sn*-glycero-3-phosphocholine (SLPC), 1-stearoyl-2-arachidonoyl-*sn*-glycero-3-phosphocholine (SAPC), 1-palmitoyl-2-docosahexaenoyl-*sn*-glycero-3-phosphocholine (PDHPC) and 1-stearoyl-2-docosahexaenoyl-*sn*-glycero-3-phosphocholine (SDHPC) and were purchased from Avanti Polar Lipids. Commercially available oxidized phospholipid standards including 1-palmitoyl-2-(5-oxovaleroyl)-*sn*-glycero-3-phosphocholine (POVPC), 1-palmitoyl-2-azelaoyl-*sn*-glycero-3-phosphocholine (PAzPC), 1-palmitoyl-2-(9-oxo)nonanoyl-*sn*-glycero-3-phosphocholine (PONPC), and 1-palmitoyl-2-glutaroyl-*sn*-glycero-3-phosphocholine (PGPC) were from Avanti Polar Lipids while 1-(palmitoyl)-2-(5-keto-6-octene-dioyl)phosphatidylcholine (KOdiA-PC) and 1-palmitoyl-2-(4-keto-dodec-3-ene-dioyl)phosphatidylcholine (KDdiA-PC) were from Cayman Chemicals. Internal standards (ISs) 2-dinonanoyl-*sn*-glycero-3-phosphocholine (DNPC) and 1-heptadecanoyl-2-docosatetraenoyl-*sn*-glycero(d5)-3-phosphocholine (17:0-22:4 PC-d5) were from Avanti Polar Lipids.

### Sample preparation of kidney tissue samples for targeted oxidized PCs analysis

Frozen kidneys were homogenized as described in “Sample preparation of kidney tissue samples for untargeted lipidomics analysis”. To 25 µL of kidney tissue homogenate (equal to 2.5 mg of frozen tissue), 20 µL of the IS mix (containing 100 ng/mL DNPC and 17:0-22:4 PC-d5 each in methanol), 500 µL LC-MS grade methanol and 500 µL chloroform were added. Samples were vortexed and 400 µL LC-MS grade water were added to the kidney tissue samples. The samples were then vortexed and centrifuged at 4000×g for 10 min and 400 µL of the lower chloroform layer were transferred to a glass vial and evaporated under a gentle stream of N_2_. Dried samples were stored at −80 °C until the day of analysis. On the day of analysis, samples were reconstituted in 50 µL 2-propanol, vortexed and sonicated for 5 min. 50 µL of LC-MS grade water were added and samples were vortexed and sonicated for 5 min again. The samples were transferred to inserts and were ready to be analyzed by the LC-MS/MS targeted oxidized PCs platform. Same as in the untargeted lipidomics analysis, a quality control (QC) was made by pooling equal aliquots of every single sample and was analyzed periodically during the analytical run.

### Targeted LC-MS/MS method for oxidized PCs

10 µL of the lipid extracts were injected in a LC-MS/MS system composed of a Shimadzu LC-40 HPLC system coupled to a Sciex QTRAP 6500+ MS system in ESI+ multiple reaction monitoring (MRM) mode. Lipid separation was achieved using a Kinetex® C18 100 Å LC Column (50 × 2.1 mm, 1.7 μm) from Phenomenex, kept at 50 °C. Mobile phase A consisted of acetonitrile:water (50:50, v/v), and mobile phase B was 2-propanol:acetonitrile:water (85:10:5, v/v/v), both containing 5 mM ammonium formate and 0.1% formic acid delivered at a flow rate of 400 µL/min. The gradient elution was 10% B at 0.0 min, 25% B at 2.0 min, 30% B at 3.5 min, 75% B at 5.0 min, 77% B at 8.5 min, 100% B at 9.0 min, 100% B at 10.0 min and reverting to 10% B at 10.5 min. Data acquisition and integration were both performed using Sciex Analyst and Sciex OS software, respectively. The peak area ratios of lipids to the ISs were used for further analysis. The MRM transitions are detailed in **Table S6**. Other MS parameters include a curtain gas of 26 psi, ion source gas 1 and 2 set to 40 and 30 psi, respectively, a “low” collision gas, an ion spray voltage of 4500 V, a source temperature of 450 °C, and an entrance, collision cell exit declustering potential of 10, 15 and 100 V, respectively.

### Oxilipidomic analysis by LC-MS/MS on perfusate

Oxylipids were analyzed from 100 µL perfusate samples according to published protocols (34). Briefly, to 100 µL perfusate was added 100 µL water, 600 µL methanol and an internal standard mix. Following acidification with formic acid, samples were cleaned up by solid phase extraction using C18 cartridges before being analyzed by LC-MS/MS in MRM mode on a Shimadzu Nexera series UHPLC system coupled to a Sciex 6500 QTrap.

### Bulk RNA-sequencing

RNA was extracted from cryopreserved kidney biopsies using RNeasy Kits for RNA purification (Qiagen, 74106), following manufacturer’s instructions. The library was constructed by BGI Laboratories using the DNBSEQ Eukaryotic Strand-specific Transcriptome Resequencing protocol. Sequencing was performed on the DNBSEQ platform with PE150 or PE100. Trimmed FastQ files were provided by BGI. After which, sequence reads were mapped to the reference genome (GRCh38/hg38) and a count matrix was generated using Rsubread (v2.20.0). Count matrices were read into DEseq2, and size factors were estimated. Read counts were analyzed using DEseq2 (v1.46.0) (35).

All bulk RNA sequencing analysis were performed using R (version 4.4.0). Gene set variation analysis (GSVA, v2.0.2) was performed for GOBP_Iron_Ion_Transport (n=58 genes; GO:0006826, 2024.1.Hs), WP_Ferroptosis (n=64 genes; WP4313, 2024.1.Hs), and FerrDb v2 validated driver, suppressor and marker genes with a protein product (n=313 genes) (18, 36).

### Statistical analysis

Statistical analysis was performed using GraphPad Prism 9. We tested for Gaussian distribution by performing the Shapiro-Wilk normality test. When normal distribution was confirmed, we used an unpaired two-tailed t-test to compare two datasets and an one-way ANOVA to compare more than two datasets. In cases where multiple comparisons involved paired observations, we used a two-way ANOVA. Bonferroni’s correction was used when multiple comparisons were made. If the normality test indicated a non-Gaussian distribution, we used the Mann-Whitney U-test to compare two datasets and a Kruskal-Wallis test to compare more than two datasets. Dunn’s correction was used when multiple comparisons were made. For the untargeted lipidomics analysis we performed unpaired t-tests with Welsch correction. Correlation coefficients, where reported, are Pearson’s Rho for normally distributed or Spearman’s rank correlation for non-normally distributed variables. A p-value of 0.05 was used as threshold for statistical significance. Data are presented as mean ± SD, unless indicated otherwise.

## Acknowledgments

We gratefully acknowledge the support provided by Rutger van Rooden (Transplant Center, LUMC, Leiden, the Netherlands) in making the 3D renderings of the organ chambers used for perfusion experiments, Mohan Ghorasaini and Kevin Brewster (Center of Proteomics and Metabolomics, LUMC, Leiden, the Netherlands) for their technical assistance with lipidomic analysis, and Manon Zuurmond (Department of Internal Medicine, LUMC, Leiden, the Netherlands) for making the illustrations.

This work is supported by the Dutch Kidney Foundation through the Participants of the Friends Lottery (20INI011). The Novo Nordisk Foundation Center for Stem Cell Medicine (reNEW) is supported by Novo Nordisk Foundation grants (NNF21CC0073729). Ton J. Rabelink is funded by the European Union through ERC grant (SPARK 101140863).

