## Supplemental material for "Normothermic human kidney preservation drives iron accumulation and ferroptosis"

Figure S1 | Perfusion dynamics during 48-hour RBC-based human kidney perfusion.
Figure S2 | Effect of cold preservation method prior to warm perfusion.
Figure S3 | Perfusion dynamics. Comparison between RBC-based perfusion with or without a deceased donor kidney in the platform.
Figure S4 | Secondary products of lipid peroxidation increase in perfusate and tissue during RBC-based human kidney perfusion.
Figure S5 | Perfusion dynamics. Comparison between dialysis-filter based free hemoglobin (fHb) removal during RBC-based perfusion with RBC-based perfusion of deceased human donor kidneys.
Figure S6 | Dialysis-based removal of free hemoglobin reduces hemolysis-driven iron accumulation and phospholipid peroxidation.
Figure S7 | Perfusion dynamics. Comparison between cell-free perfusion with RBC-based perfusion of deceased human donor kidneys.
Figure S8 | Effect of donor background on cell-free perfusion.
Figure S9 | Relation between donor final serum creatinine and kidney-specific injury markers during perfusion.

Table S1 | Comparison of deceased donor characteristics between the different perfusion groups.
Table S2 | Perfusate and substitution solution composition.
Table S3 | Tissue histology scoring.
Table S4 | Overview of lipid (sub-)classes identified through untargeted lipidomics.
Table S5 | Tissue lipid species changes following RBC-based normothermic kidney perfusion.
Table S6 | Multiple reaction monitoring transitions used for the identification of the different PCs and oxidized PCs.

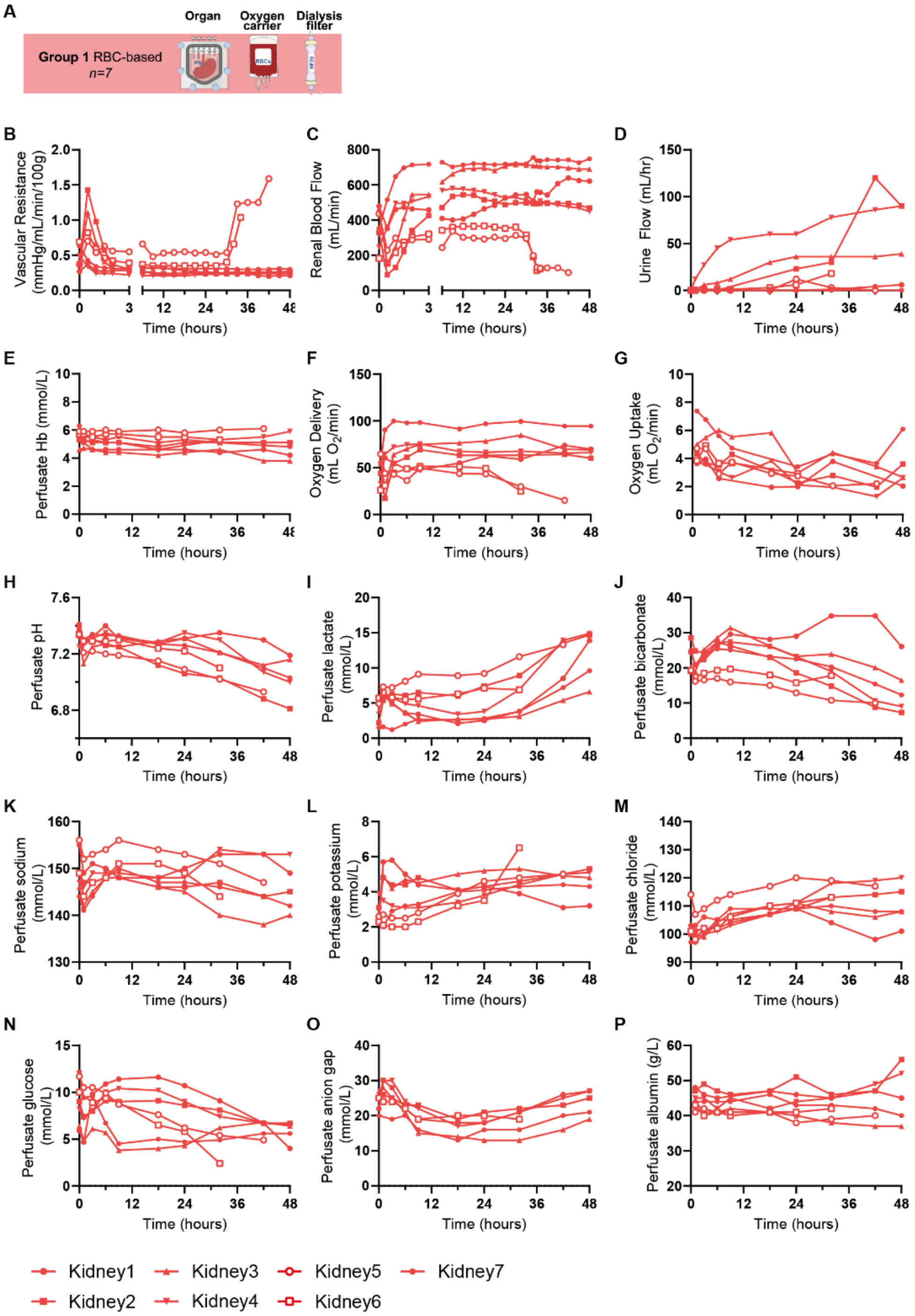

**Figure S1 related to Figure 1. Perfusion dynamics during 48-hour RBC-based human kidney perfusion. A**, Schematic overview illustrating the experimental conditions. **B**, Vascular resistance. During ex situ reperfusion an early increase in vascular resistance (and decrease in renal blood flow) was observed. Within 1-2 hours this restored to a steady-state that was maintained for 48-hours in 5 of the 7 perfused human kidneys. **C**, Renal blood flow. **D**, Urine flow. **E**, Perfusate Hemoglobin (Hb). **F**, Oxygen delivery. **G**, Oxygen uptake. **H**, Perfusate pH. **I**, Perfusate lactate. **J**, Perfusate bicarbonate. **K**, Perfusate sodium. **L**, Perfusate potassium. **M**, Perfusate chloride. **N**, Perfusate glucose. **O**, Perfusate anion gap. **P**, Perfusate albumin.

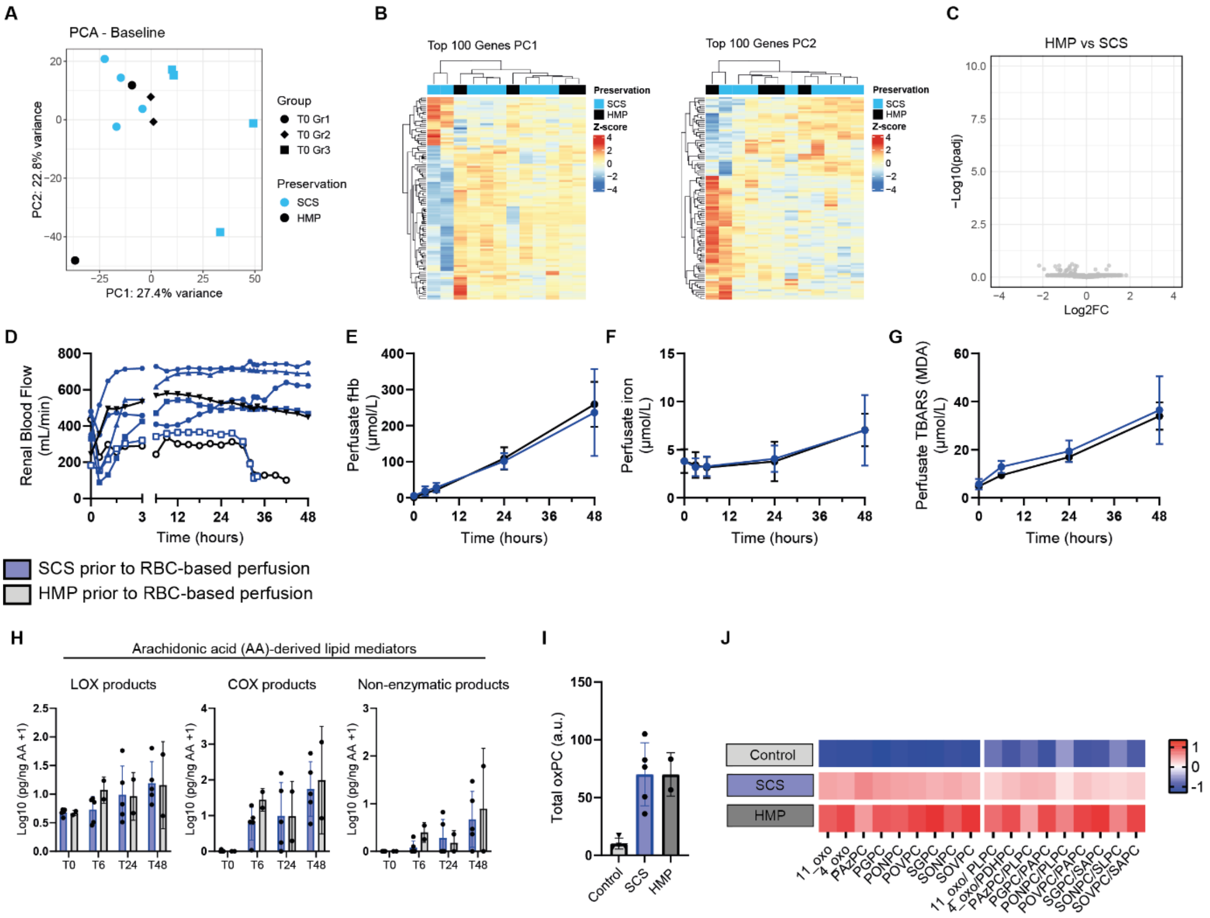

**Figure S2 related to Figure 1. Effect of cold preservation method prior to warm perfusion.**

**A-C**, Transcriptomics on human kidneys that were cold preserved on either HMP or SCS. PCA illustrates the variance between samples (**A**), with heatmap of the Top100 genes belonging to PC1 and PC2 illustrating that samples do not cluster based on their transcriptome (**B**). Volcano plot illustrates the absence of DEGs between HMP and SCS preserved human kidneys (**C**).

D-J, No differences were observed in RBC-perfused kidneys that were either preserved on SCS or HMP, looking at renal blood flow (**D**), hemolysis (**E,F**), and perfusate TBARS (**G**), perfusate lipid mediators (**H**), and tissue oxPCs (**I,J**).

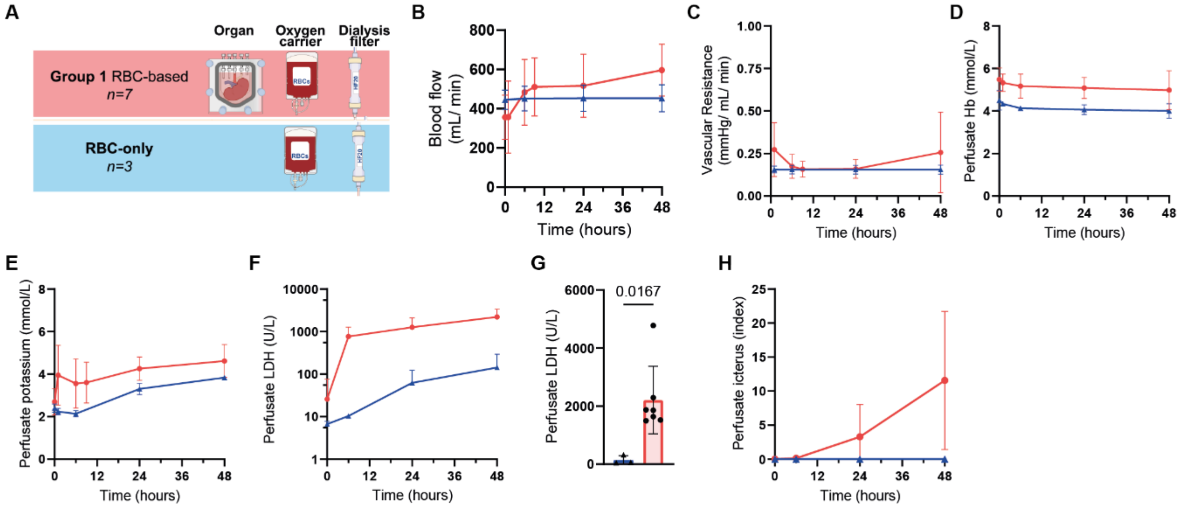

**Figure S3 related to Figure 1. Perfusion dynamics.** Comparison between RBC-based perfusion with (n=7) or without (n=3) a deceased donor kidney in the platform. **A**, Schematic overview illustrating the experimental conditions. **B**, Blood flow. **C**, Vascular resistance. **D**, Perfusate Hb. **E**, Perfusate potassium. **F**, Perfusate LDH. **G**, Perfusate LDH after 48-hour perfusion (Mann-Whitney test). **H**, Perfusate icterus index. Indicator for amount of total bilirubin in perfusate.

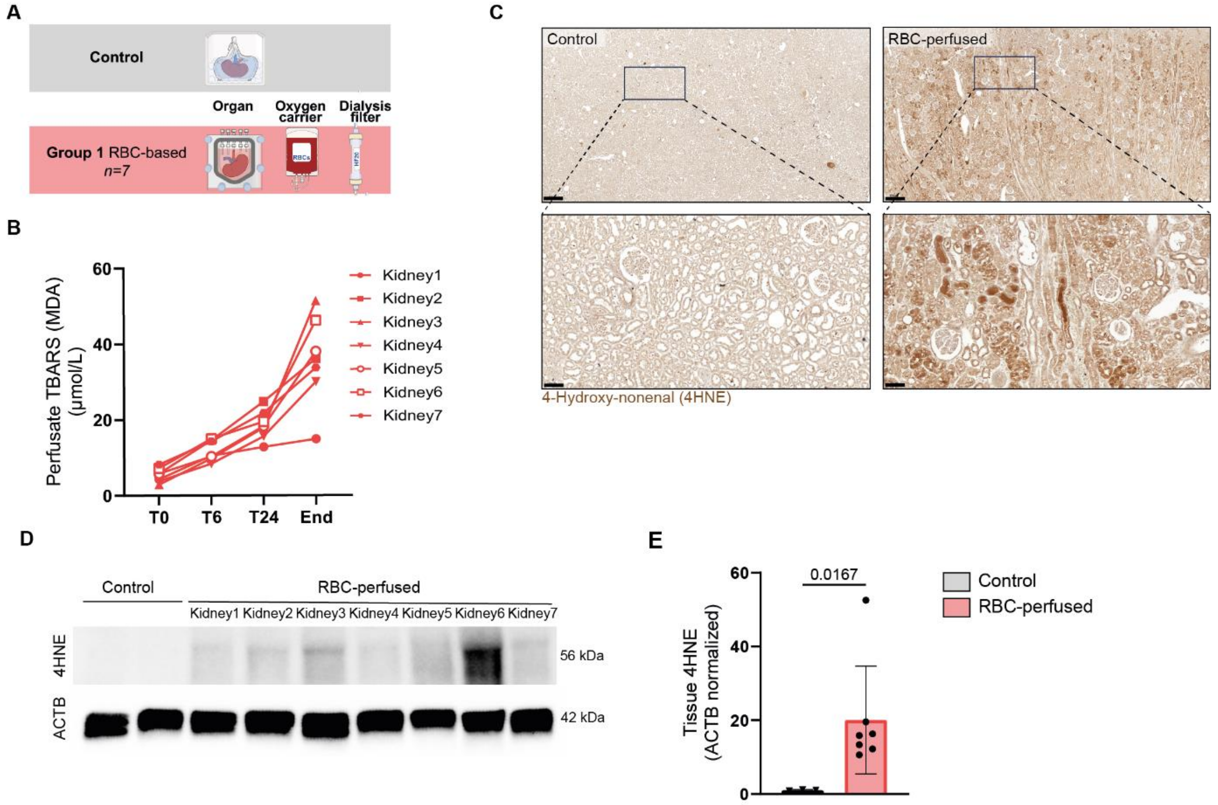

**Figure S4 related to Figure 2. Secondary products of lipid peroxidation increase in perfusate and tissue during RBC-based human kidney perfusion. A**, Schematic overview illustrating the experimental conditions. **B**, Perfusate TBARS (MDA) (Repeated measures one-way ANOVA). **C**, Immunohistochemistry for 4-hydroxynonenal (4HNE). Representative images of control (n=3) and RBC-perfused human kidneys (n=7). RBC-perfused Kidney3 is shown. Scale bars represent 500 µm (top panels) and 100 µm (bottom panels). **D**, Tissue 4HNE abundance in control and RBC-perfused human kidneys. B-actin (ACTB) for normalization. **E**, Densitometric quantification. Tissue 4HNE levels in RBC-perfused human kidneys relative to control kidney biopsies. Normalized to ACTB levels (Mann-Whitney test).

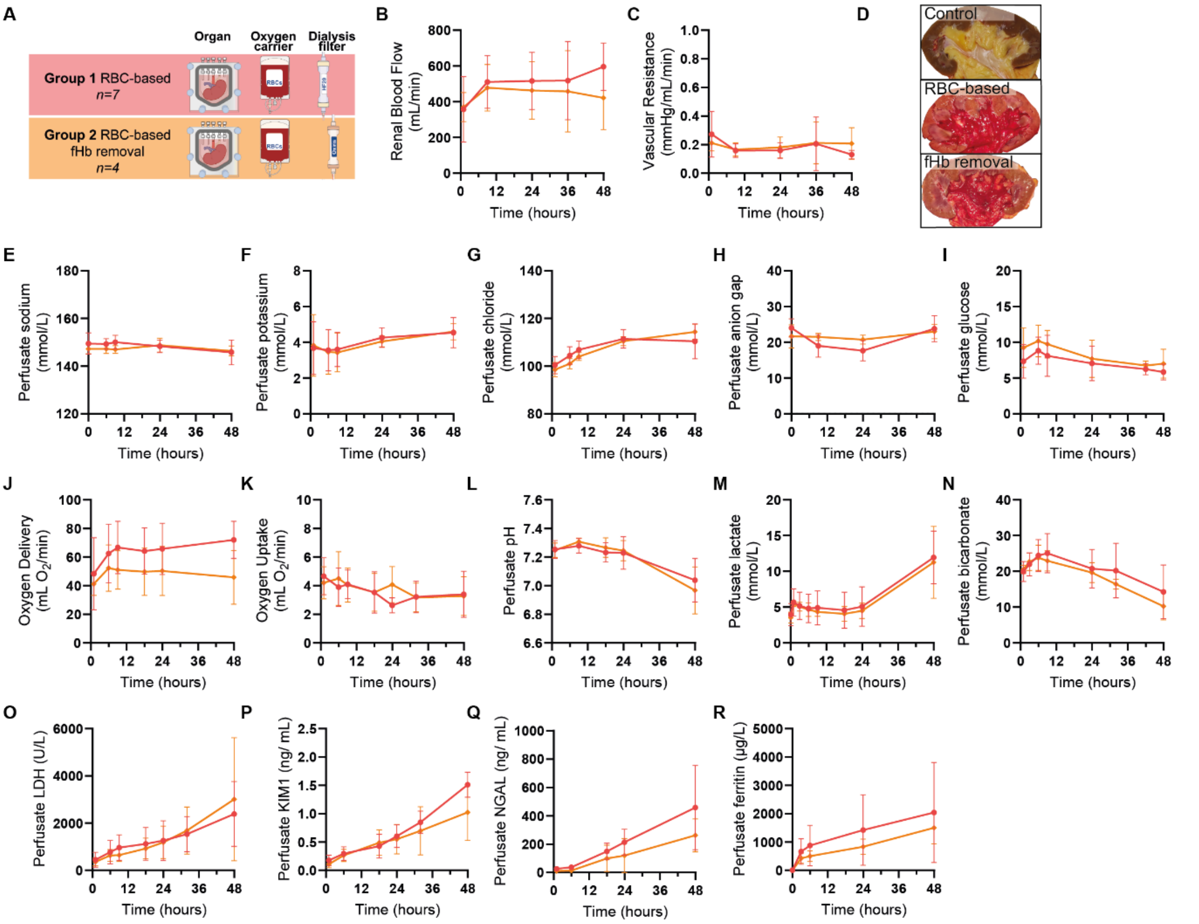

**Figure S5 related to Figure 3. Perfusion dynamics.** Comparison between dialysis-filter based free hemoglobin (fHb) removal during RBC-based perfusion (Group2, n=4) with RBC-based perfusion (Group1, n=7) of deceased human donor kidneys. **A**, Schematic overview illustrating the experimental conditions. **B**, Renal blood flow. **C**, Vascular resistance. **D**, Macroscopic appearance. **E**, Perfusate sodium. **F**, Perfusate potassium. **G**, Perfusate chloride. **H**, Perfusate anion gap. **I**, Perfusate glucose. **J**, Oxygen delivery. **K**, Oxygen uptake. **L**, Perfusate pH. **M**, Perfusate lactate. **N**, Perfusate bicarbonate. **O**, Perfusate LDH. **P**, Perfusate KIM1. **Q**, Perfusate NGAL. **R**, Perfusate ferritin. Data are presented as mean with SD.

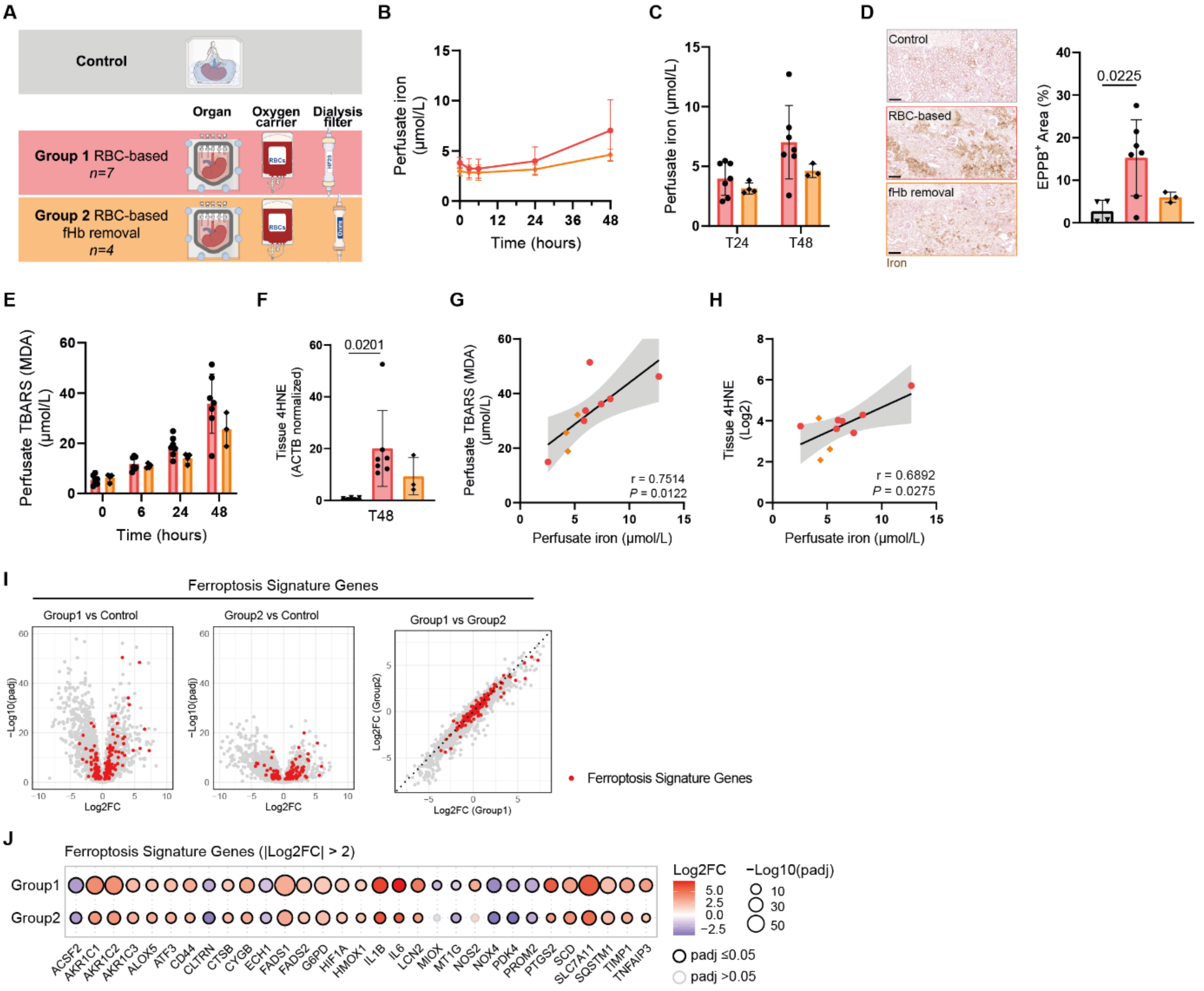

**Figure S6 related to Figure 3. Dialysis-based removal of free hemoglobin reduces hemolysis-driven iron accumulation and phospholipid peroxidation. A**, Schematic overview illustrating the experimental conditions. **B-C**, Perfusate iron during perfusion (**B**) and perfusate iron at 24 and 48 hour timepoints (**C**) (Two-way ANOVA). **D**, Enhanced Perl's Prussian blue (EPPB) staining for iron deposition in tissue. Representative images and quantification (One-way ANOVA). Scale bar, 200 µm. **E**,Perfusate TBARS (MDA). **F**, Densitometric quantification of 4HNE levels in tissue at the end of perfusion, relative to control kidney biopsies. Normalized to ACTB (Kruskal-Wallis test). **G**, Association between perfusate iron and perfusate TBARS (MDA) after 48 hour perfusion (Pearson's correlation coefficient). **H**, Association between perfusate iron and tissue 4-hydroxynonenal (4HNE) after 48 hour perfusion (Pearson's correlation coefficient).

**I**, Volcano plots depicting significantly up- and downregulated genes (Log2FC and -Log10(padj)) in kidneys after RBC-based perfusion or fHb removal during RBC-based perfusion, compared to control kidneys (Left and middle panel). The right panel shows the correlation between Log2FC for RBC-perfused vs control and fHb removal vs control kidneys. Red highlights validated FerrDb v2 driver, suppressor, or marker genes with a protein product (n=313). **J**, Dotplot illustrating dynamics in FerrDb v2 driver, suppressor, and marker genes after RBC-perfused and fHb removal conditions, compared to control kidneys. Genes included have |Log2FC|≥2 and padj ≤ 0.05 for at least one of the comparisons.

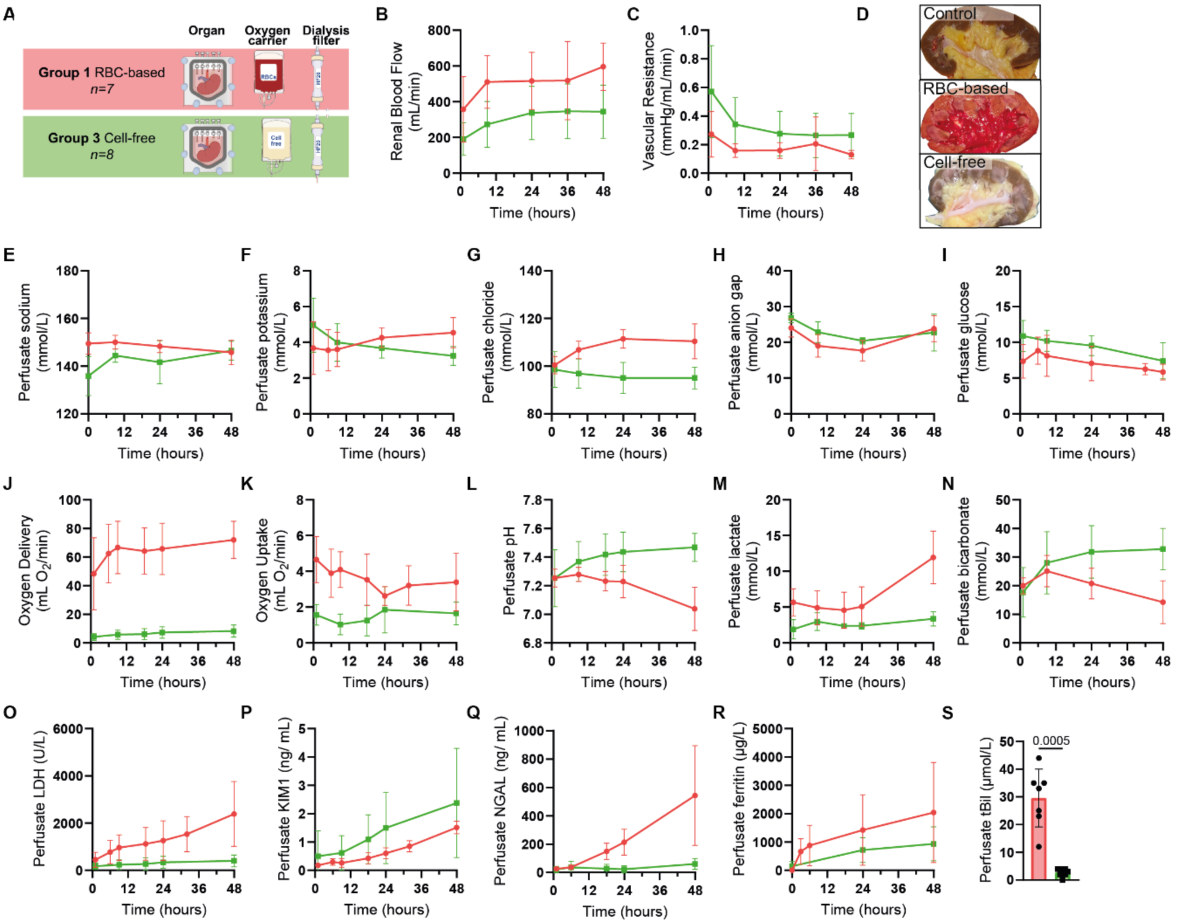

**Figure S7 related to Figure 4. Perfusion dynamics.** Comparison between cell-free perfusion (Group3, n=8) with RBC-based perfusion (Group1, n=7) of deceased human donor kidneys. **A**, Schematic overview illustrating the experimental conditions. **B**, Renal blood flow. **C**, Vascular resistance. **D**, Macroscopic appearance. **E**, Perfusate sodium. **F**, Perfusate potassium. **G**, Perfusate chloride. **H**, Perfusate anion gap. **I**, Perfusate glucose. **J**, Oxygen delivery. **K**, Oxygen uptake. **L**, Perfusate pH. **M**, Perfusate lactate. **N**, Perfusate bicarbonate. **O**, Perfusate LDH. **P**, Perfusate KIM1. **Q**, Perfusate NGAL. **R**, Perfusate ferritin. **S**, Perfusate total bilirubin after 48 hour perfusion (Unpaired t-test with Welch's correction). Data are presented as mean with SD.

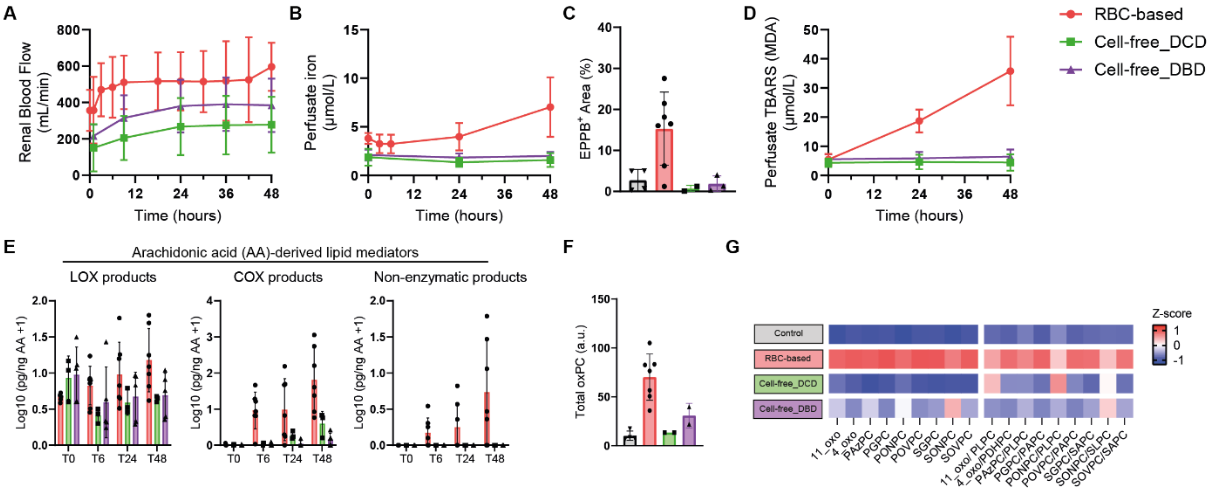

**Figure S8 related to Figure 4. Donor background prior to cell-free perfusion.** The RBC-based group (Group1) solely contained DCD donors (n=7). The cell-free perfusion group (Group3) contained both DCD and DBD donors (n=3 and 5, respectively). We found donor background has a negligible impact on perfusion outcomes. **A**, Renal blood flow. **B**, Perfusate iron. **C**, Tissue iron based on EPPB histology. **D**, Perfusate TBARS. **E**, Perfusate oxilipidomics. **F**, Tissue total oxPCs. **G**, Tissue oxPC.

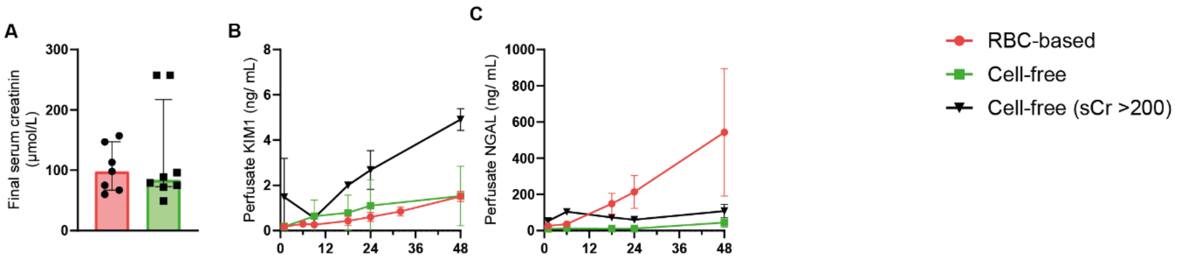

**Figure S9 related to Figure 4. Relation between donor final serum creatinine and kidney-specific injury markers during perfusion.** **A**, Final serum creatinine. **B**, Perfusate KIM1. **C**, Perfusate NGAL.

**Table S1 | Comparison of deceased donor characteristics between the different perfusion groups.** Group1 consists of kidneys perfused according to the initial protocol, which involved stored red blood cells (RBCs) and an HF20 dialysis filter. Group2 kidneys were perfused with stored RBCs and an Oxiris dialysis filter, allowing the removal of free hemoglobin (fHb). Group3 kidneys were perfused without stored RBCs (i.e. in the absence of fHb) at subnormothermia (25°C).

| | **Group1** (n=7) | **Group2** (n=4) | **Group3** (n=8) |
|---|---|---|---|
| **Donor type** (DCD)[a] | 7 (100%) | 4 (100%) | 3 (38%) |
| **Age** (years)[b] | 58±13 | 68±9 | 61±11 |
| **Sex** (male)[a] | 7 (100%) | 3 (75%) | 6 (75%) |
| **BMI** (kg/m$^2$)[b] | 26±4 | 27±2 | 24±5 |
| **ICU stay** (days)[c] | 2 (1-3) | 2 (2-7) | 1.5 (1-3) |
| **Final serum creatinine** (µmol/L)[c] | 98 (67-147) | 80 (53-187) | 84 (73-217) |
| **Cause of death**[a] | | | |
| *- Circulational: cardiac arrest* | 1 (14%) | 1 (25%) | 3 (38%) |
| *- CVA: cerebral ischemia* | 2 (29%) | 1 (25%) | 1 (13%) |
| *- CVA: intra cerebral bleeding* | 1 (14%) | 0 | 2 (25%) |
| *- Suicide: respiratory* | 2 (29%) | 0 | 0 |
| *- Suicide: jump* | 1 (14%) | 0 | 0 |
| *- SDH: Sub Dural Hematoma* | 0 | 1 (25%) | 0 |
| *- SAB: Sub Arachnoidal Bleeding* | 0 | 1 (25%) | 1 (13%) |
| *- Euthanasia* | 0 | 0 | 1 (13%) |
| **Cold preservation method**[a] | | | |
| *- SCS* | 5 (71%) | 1 (25%) | 7 (87%) |
| *- HMP* | 2 (29%) | 3 (75%) | 1 (13%) |
| **Cold preservation time** (hours) | 20 (18-23) | 24 (21-26) | 15 (12-18)[¥] |
| **Stored RBC age** (days)[c] | 16 (2-23) | 25 (13-33) | *NA* |

Data are n (%) for categorical variables ([a]) and mean ± SD ([b]) or median (IQR) ([c]) for continuous variables. [¥] The exact cold preservation time (in hours) is unknown for 3 of 8 kidneys, but cell-free perfusion began within 24 hours of procurement.

**Table S2 | Perfusate and substitution solution composition.**

| **Perfusate composition** | | | | |
|---|---|---|---|---|
| | **Stock [c]** | **Per 1L** | **Final[c]** | **Comments** |
| Washed pRBCs[1] | Hct 50-60% | 500 mL | Hct 20-25% | Sanquin |
| DMEM F12, HEPES | N/A | 200 mL | N/A | Cat# 11330, Gibco. |
| Human Serum Albumin | 200 gr $L^{-1}$ | 200 mL | 40 gr $L^{-1}$ | Alburex 20, CSL Behring bv. |
| Sodium Hydroxide | 1M | 12.5 mL | 12.5 mmol $L^{-1}$ | Cat# 567530, Calbiochem, Merck. |
| Sterile water | N/A | 75 mL | N/A | Versylene Fresenius. |
| Sodium bicarbonate | 7.5% | 10 mL | 0.075% | Cat# 25080094, Gibco. |
| Creatinine | 50 mg $mL^{-1}$ | 1 mL | 50 mg $L^{-1}$ | C4255-25G, Sigma-Aldrich |
| Heparin | 5000 U $mL^{-1}$ | 0.2 mL | 1000 U $L^{-1}$ | LEO. |
| Penicillin-streptomycin | 5000 U $mL^{-1}$<br>5000 µg $mL^{-1}$ | 10mL | 50 U $mL^{-1}$<br>50 µg $mL^{-1}$ | Cat# 15070063, Gibco. |
| Ciprofloxacin | 2 mg $mL^{-1}$ | 3 mL | 6 µg $mL^{-1}$ | Fresenius Kabi. |
| Fungizone | 0.25 mg $mL^{-1}$ | 1 mL | 0.25 µg $mL^{-1}$ | Bristol-Myers Squibb. |
| Insulin | 1 U $mL^{-1}$ | 1 mL | 1U $L^{-1}$ | Solostar, Lantus (100 U $mL^{-1}$) |
| **Substitution solution composition** | | | | |
| | **Stock [c]** | **Per 1L** | **Final[c]** | **Comments** |
| DMEM F12, HEPES | N/A | 940 mL | N/A | Cat# 11330, Gibco. |
| Sodium Hydroxide | 1M | 6 mL | 12.5 mmol $L^{-1}$ | Cat# 567530, Calbiochem, Merck. |
| Sterile water | N/A | 30 mL | N/A | Versylene Fresenius. |
| Sodium bicarbonate | 7.5% | 10 mL | 0.075% | Cat# 25080094, Gibco. |
| Creatinine | 50 mg $mL^{-1}$ | 1 mL | 50 mg $L^{-1}$ | C4255-25G, Sigma-Aldrich |
| Heparin | 5000 U $mL^{-1}$ | 0.2 mL | 1000 U $L^{-1}$ | LEO. |
| Penicillin-streptomycin | 5000 U $mL^{-1}$<br>5000 µg $mL^{-1}$ | 10mL | 50 U $mL^{-1}$<br>50 µg $mL^{-1}$ | Cat# 15070063, Gibco. |
| Ciprofloxacin | 2 mg $mL^{-1}$ | 3 mL | 6 µg $mL^{-1}$ | Fresenius Kabi. |
| Fungizone | 0.25 mg $mL^{-1}$ | 1 mL | 0.25 µg $mL^{-1}$ | Bristol-Myers Squibb. |
| Insulin | 1 U $mL^{-1}$ | 1 mL | 1U $L^{-1}$ | Solostar, Lantus (100 U $mL^{-1}$) |

[1] Prior to their use, RBCs (pRBCs) were washed three times. Briefly, RBCs were transferred from blood bags to 250 mL conicles (±166 mL pRBCs/conicle), supplemented with DMEM F12 to 250 mL and centrifuged at 1000g for 10 minutes (21°C, acc9, dec4). After centrifugation the supernatant was removed. This was repeated three times. Washed RBCs were added to the perfusate once the perfusion platform had been primed with the other perfusate components.

**Table S3 | Tissue histology scoring.**

| **I. Histological scores over the course of RBC-based human kidney perfusion (*paired samples*)** | | | | | | | |
|---|---|---|---|---|---|---|---|
| | Baseline *(n=7)* | T6 *(n=7)* | $P^{*}$ | T24 *(n=7)* | $P^{*}$ | T48 *(n=7)* | $P^{*}$ |
| Acute Tubular Injury [0-3][a] | 2 [1-2] | 1 [1-2] | 0.29 | 1 [1-2] | >0.99 | 1 [1-2] | >0.99 |
| Tubular casts [0-3][b] | 2 [1-2] | 1 [0-2] | 0.64 | 0 [0-1] | 0.09 | 1 [0-3] | 0.36 |
| Tubular vacuolization [0-3][b] | 1 [1-2] | 2 [1-3] | 0.24 | **3 [3-3]** | **0.00** | **3 [1-3]** | **0.04** |
| Interstitial edema [0-3][b] | 2 [1-3] | 1 [0-2] | 0.29 | 1 [0-3] | 0.64 | 2 [1-3] | >0.99 |
| Tubular dilation [0-3][b] | 2 [2-3] | 2 [1-3] | >0.99 | **1 [0-2]** | **0.01** | **1 [0-2]** | **0.07** |
| **II. Histological scores at the end of the different perfusion modalities compared to baseline** | | | | | | | |
| | Baseline *(n=14)* | Group1 *(n=7)* | $P^{¥}$ | Group2 *(n=3)* | $P^{¥}$ | Group3 *(n=8)* | $P^{¥}$ |
| Acute Tubular Injury [0-3][a] | 1.5 [1-2] | 1 [1-2] | >0.99 | 1 [1-2] | >0.99 | 1.5 [1-2] | >0.99 |
| Tubular casts [0-3][b] | 1.5 [0-3] | 1 [0-3] | >0.99 | 2 [1-2] | >0.99 | 1 [1-3] | >0.99 |
| Tubular vacuolization [0-3][b] | 1 [0-3] | **3 [1-3]** | **0.00** | **3 [3-3]** | **0.01** | **2 [2-3]** | **0.02** |
| Interstitial edema [0-3][b] | 1.5 [1-3] | 2 [1-3] | >0.99 | 3 [2-3] | 0.19 | 2.5 [2-3] | 0.09 |
| Tubular dilation [0-3][b] | 2 [1-3] | **1 [0-2]** | **0.04** | **1 [0-1]** | **0.03** | 1.5 [0-3] | 0.38 |

Data are Median (Min-Max). [a] 0, Absent; 1, Loss of brush border/ vacuolization of tubules; 2, Cell detachment/ cellular casts; 3, Necrosis. [b] 0, 0-1%; 1, >1-10%; 2, >10-25%; 3, >25%. Group1, RBC-based perfusion; Group2, dialysis-based fHb removal during RBC-based perfusion; Group3, Cell-free perfusion.

[*] *P*-value compares mean rank of Timepoint with Baseline based on Friedman test with Dunns correction for multiple comparison.

[¥] *P*-value compares mean rank of Timepoint with Baseline based on Friedman test with Dunns correction for multiple comparison.

**Table S4 | Overview of lipid (sub-)classes identified through untargeted lipidomics.** Lipid species were classified according to the MS-DIAL nomenclature.

| Categories | # | Main class | # | Lipid subclass | Abbreviation | # |
|---|---|---|---|---|---|---|
| Fatt Acyls [FA] | 53 | Fatty acids and Conjugates | 23 | Free fatty acid | FA | 23 |
| | | Fatty esters | 30 | Acylcarnitine | CAR | 30 |
| Glycerolipids [GL] | 186 | Diradylglycerols | 36 | Diacylglycerol | DG | 35 |
| | | | | Ether-linked diacylglycerol | EtherDG | 1 |
| | | Triradylglycerols | 150 | Triacylglycerol | TG | 125 |
| | | | | Oxidized triglyceride | OxTG | 16 |
| | | | | Ether-linked triacylglycerol | EtherTG | 9 |
| Glycerophospholipids [GP] | 300 | Phospholipids | 167 | Phosphatidylcholine | PC | 38 |
| | | | | Phosphatidylethanolamine | PE | 51 |
| | | | | Phosphatidylserine | PS | 21 |
| | | | | Phosphatidylglycerol | PG | 36 |
| | | | | Phosphatidylinositol | PI | 18 |
| | | | | Phosphatidic acid | PA | 3 |
| | | Lyso Phospholipids | 44 | Lysophophatidylcholine | LPC | 16 |
| | | | | Lysophosphatidylethanolamine | LPE | 23 |
| | | | | Lysophosphatidylserine | LPS | 2 |
| | | | | Lysophosphatidylinositol | LPI | 3 |
| | | Ether-linked Phospholipids | 51 | Ether-linked phosphatidylcholine | EtherPC | 13 |
| | | | | Ether-linked phosphatidylethanolamine | EtherPE | 30 |
| | | | | Ether-linked lysophosphatidylethanolamine | EtherLPE | 6 |
| | | | | Ether-linked phosphatidylinositol | EtherPI | 2 |
| | | Oxidized Phospholipids | 22 | Oxidized phosphatidylcholine | OxPC | 18 |
| | | | | Oxidized phosphatidylinositol | OxPI | 3 |
| | | | | Ether-linked oxidized phosphatidylethanolamine | EtherOxPE | 1 |
| | | Other-Phospholipids | 16 | Bismonoacylglycerophosphate | BMP | 6 |
| | | | | Cardiolipin | CL | 9 |
| | | | | Lysocardiolipin | MLCL | 1 |
| Sphingolipids [SP] | 113 | Ceramides [SP02] | 37 | Ceramide alpha-hydroxy fatty acid-phytospingosine | Cer_AP | 5 |
| | | | | Ceramide alpha-hydroxy fatty acid-sphingosine | Cer_AS | 1 |
| | | | | Ceramide non-hydroxyfatty acid-dihydrosphingosine | Cer_NDS | 4 |
| | | | | Ceramide non-hydroxyfatty acid-phytospingosine | Cer_NP | 9 |
| | | | | Ceramide non-hydroxyfatty acid-sphingosine | Cer_NS | 18 |
| | | Neutral glycosphingolipids [SP05] | 30 | Hexosylceramide alpha-hydroxy fatty acid-phytospingosine | HexCer_AP | 4 |
| | | | | Hexosylceramide hydroxyfatty acid-dihydrosphingosine | HexCer_HDS | 3 |
| | | | | Hexosylceramide non-hydroxyfatty acid-sphingosine | HexCer_NS | 4 |
| | | | | Dihexosylceramide | Hex2Cer | 8 |
| | | | | Trihexosylceramide | Hex3Cer | 11 |
| | | Acidic glycosphingolipids [SP06] | 6 | Sulfatide | SHexCer | 6 |
| | | Phosphosphingolipids [SP03] | 40 | Sphingomyelin | SM | 30 |
| | | | | Ceramide phosphoethanolamine | PE-Cer | 1 |
| | | | | Ceramide phosphoinositol | PI_Cer | 7 |
| | | | | Oxidized ceramide phosphoinositol | Ox_PI_Cer | 2 |
| Sterol Lipids [ST] | 10 | Sterols [ST01] | 8 | Cholesteryl ester | CE | 8 |
| | | Steroid conjugates [ST05] | 2 | Sterol sulfate | Ssulfate | 2 |

**Table S5 | Tissue lipid species changes following RBC-based normothermic kidney perfusion as defined by Log2FC ≥1.5 and *P* value <0.05.** Based on total area normalized lipid species as measured through untargeted lipidomics in perfused RBC-based kidney biopsies (n=7) in comparison to unperfused deceased donor kidney biopsies (n=3).

| Lipid class | Lipid name | Fold change (Perfused/ Unperfused) | *P* value |
|---|---|---|---|
| **OxPC** | PC 32:2;O2 | 7.536 | 0.0104 |
| **OxPC** | PC 30:1;O2 | 7.007 | 0.0197 |
| **EtherPC** | PC O-28:0 | 6.098 | 0.0002 |
| **OxPI** | PI 32:1;O2 | 4.973 | 0.0046 |
| **EtherPE** | PE O-32:4_2 | 4.355 | 0.0013 |
| **OxPC** | PC 34:1;O2 | 4.018 | 0.0320 |
| **OxPC** | PC 36:3;O3 | 3.393 | 0.0181 |
| **FA** | FA 26:5 | 3.164 | 0.0022 |
| **HexCer_NS** | HexCer 42:3;O2_2 | 2.631 | 0.0009 |
| **OxPC** | PC 36:2;O2 | 2.553 | 0.0030 |
| **PC** | PC 42:2_1 | 2.484 | 0.0010 |
| **OxTG** | TG 46:1;O2 | 2.465 | 0.0185 |
| **HexCer_NS** | HexCer 42:2;O2_2 | 2.456 | 0.0016 |
| **EtherOxPE** | PE O-36:5;O3 | 2.452 | 0.0060 |
| **PG** | PG 32:2_2 | 2.435 | 0.0360 |
| **OxPC** | PC 34:3;O_1 | 2.429 | 0.0053 |
| **OxPC** | PC 36:3;O2 | 2.386 | 0.0058 |
| **OxPI** | PI 38:5;O_1 | 2.383 | 0.0112 |
| **PC** | PC 28:0_1 | 2.381 | 0.0136 |
| **PC** | PC 42:1_2 | 2.336 | 0.0004 |
| **OxPI** | PI 38:5;O_2 | 2.308 | 0.0049 |
| **PC** | PC 29:0_1 | 2.253 | 0.0230 |
| **OxPC** | PC 34:3;O2_1 | 2.251 | 0.0020 |
| **OxPC** | PC 34:3;O_2 | 2.197 | 0.0065 |
| **PI_Cer** | PI-Cer 38:4;O3 | 2.196 | 0.0101 |
| **OxPC** | PC 36:4;O2_1 | 2.153 | 0.0058 |
| **OxPC** | PC 34:2;O2_1 | 2.123 | 0.0082 |
| **PC** | PC 32:1_2 | 1.988 | 0.0419 |
| **OxPC** | PC 36:5;O3_2 | 1.901 | 0.0105 |
| **PG** | PG 36:2 | 1.890 | 0.0150 |
| **PE** | PE 34:1_1 | 1.886 | 0.0259 |
| **OxPC** | PC 36:5;O3_1 | 1.875 | 0.0190 |
| **PI_Cer** | PI-Cer 41:8;O3 | 1.808 | 0.0114 |
| **PG** | PG 38:3 | 1.744 | 0.0128 |
| **OxPC** | PC 34:3;O2_2 | 1.707 | 0.0163 |
| **PG** | PG 32:0 | 1.701 | 0.0104 |
| **PI** | PI 40:6_2 | 1.633 | 0.0487 |
| **PE** | PE 33:1_1 | 1.630 | 0.0256 |
| **PE** | PE 40:1_2 | 1.629 | 0.0009 |
| **PE** | PE 42:1_2 | 1.602 | 0.0015 |
| **PI** | PI 40:5_3 | 1.593 | 0.0085 |
| **PE** | PE 42:2_1 | 1.584 | 0.0069 |
| **PE** | PE 40:3 | 1.576 | 0.0043 |
| **PI** | PI 40:4_3 | 1.568 | 0.0007 |
| **Cer_NS** | Cer 38:2;O2_1 | 1.565 | 0.0208 |
| **PG** | PG 36:1 | 1.563 | 0.0050 |
| **PE** | PE 37:1 | 1.544 | 0.0080 |
| **PG** | PG 40:5 | 1.541 | 0.0155 |
| **PC** | PC 30:0_1 | 1.539 | 0.0104 |
| **LPI** | LPI 18:1 | 1.528 | 0.0471 |

**Table S6 | Multiple reaction monitoring (MRM) transitions used for the identification of the different PCs and oxidized PCs and the respective internal standards (ISs).**

| Analyte | IS used for correction | Retention time (min) | Experimental MRM transition (Q1 → Q3, *m/z*) | Collision energy (V) |
|---|---|---|---|---|
| 11-oxo-9-undecenoyl-PPC | DNPC | 4.9 | 676.6 → 184.0 | 37 |
| 4-oxo-butyryl-PC | DNPC | 3.0 | 580.3 → 184.0 | 39 |
| HODA SPC | 17:0 22:4 PC-d5 | 6.7 | 734.3 → 184.0 | 42 |
| KDiAPC | DNPC | 3.8 | 720.6 → 184.0 | 40 |
| KODA PPC | DNPC | 4.9 | 704.6 → 184.0 | 40 |
| KOdiA-PC | DNPC | 2.8 | 664.6 → 184.0 | 39 |
| LPC 16:0 | DNPC | 2.5 | 496.3 → 184.0 | 33 |
| PAPC | 17:0 22:4 PC-d5 | 6.6 | 782.5 → 184.0 | 40 |
| PAzPC | DNPC | 4.0 | 666.6 → 184.0 | 40 |
| PDHPC | 17:0 22:4 PC-d5 | 6.5 | 806.5 → 184.0 | 41 |
| PDHPC-OOH | 17:0 22:4 PC-d5 | 6.0 | 838.4 → 184.0 | 41 |
| PGPC | DNPC | 2.9 | 610.6 → 184.0 | 35 |
| PLPC | 17:0 22:4 PC-d5 | 6.6 | 758.4 → 184.0 | 41 |
| PLPC OH | 17:0 22:4 PC-d5 | 6.0 | 774.6 → 184.0 | 42 |
| PLPC OOH | 17:0 22:4 PC-d5 | 6.0 | 790.6 → 184.0 | 36 |
| PONPC | DNPC | 4.8 | 650.6 → 184.0 | 34 |
| POVPC | DNPC | 3.2 | 594.6 → 184.0 | 35 |
| SAPC | 17:0 22:4 PC-d5 | 6.8 | 810.6 → 184.0 | 41 |
| SAPC OH | 17:0 22:4 PC-d5 | 6.5 | 826.4 → 184.0 | 45 |
| SAPC OOH | 17:0 22:4 PC-d5 | 6.4 | 842.4 → 184.0 | 43 |
| SAzPC | DNPC | 5.3 | 694.3 → 184.0 | 42 |
| SDHPC | 17:0 22:4 PC-d5 | 6.7 | 834.6 → 184.0 | 41 |
| SDHPC OOH | 17:0 22:4 PC-d5 | 6.3 | 866.5 → 184.0 | 39 |
| SGPC | DNPC | 4.5 | 638.2 → 184.0 | 37 |
| SLPC | 17:0 22:4 PC-d5 | 6.9 | 786.2 → 184.0 | 39 |
| SLPC epoxy keto | 17:0 22:4 PC-d5 | 6.1 | 816.4 → 184.0 | 40 |
| SLPC keto | 17:0 22:4 PC-d5 | 7.0 | 800.4 → 184.0 | 40 |
| SLPC OH | 17:0 22:4 PC-d5 | 6.4 | 802.4 → 184.0 | 38 |
| SLPC OOH | 17:0 22:4 PC-d5 | 6.2 | 818.4 → 184.0 | 36 |
| SONPC | DNPC | 5.6 | 678.3 → 184.0 | 38 |
| SOVPC | DNPC | 4.9 | 622.4 → 184.0 | 35 |
| DNPC (IS) | - | 2.4 | 538.7 → 184.0 | 46 |
| 17:0 22:4 PC-d5 (IS) | - | 6.9 | 830.2 → 184.0 | 46 |

Abbreviations: HODA SPC: 1-stearoyl-2-(9-hydroxy-12-oxododec-10-enoyl)-sn-glycero-3-phosphocholine; KDiAPC: 1-palmitoyl-2-(4-keto-dodec-3-ene-dioyl)-sn-glycero-3-phosphocholine; KODA PPC: 1-palmitoyl-2-(9,12-dioxododec-10-enoyl)-sn-glycero-3-phosphocholine; KOdiA-PC: 1-(palmitoyl)-2-(5-keto-6-octene-dioyl)- sn-glycero-3-phosphocholine; LPC 16:0: 1-palmitoyl-2-hydroxy-sn-glycero-3-phosphocholine; OH: hydroxy; OOH: hydroperoxyl; PAPC: 1-palmitoyl-2-arachidonoyl-sn-glycero-3-phosphocholine; PAzPC: 1-palmitoyl-2-azelaoyl-sn-glycero-3-phosphocholine; PDHPC: 1-palmitoyl-2-docosahexaenoyl-sn-glycero-3-phosphocholine; PGPC: 1-palmitoyl-2-glutaroyl-sn-glycero-3-phosphocholine; PLPC: 1-palmitoyl-2-linoleoyl-sn-glycero-3-phosphocholine; PONPC: 1-palmitoyl-2-(9-oxo)nonanoyl-sn-glycero-3-phosphocholine; POVPC: 1-palmitoyl-2-(5-oxovaleroyl)-sn-glycero-3-phosphocholine; SAPC: 1-stearoyl-2-arachidonoyl-sn-glycero-3-phosphocholine; SAzPCL 1-stearoyl-2-azelaoyl-sn-glycero-3-phosphocholine; SDHPC: 1-stearoyl-2-docosahexaenoyl-sn-glycero-3-phosphocholine; SGPC: 1-stearoyl-2-glutaroyl-sn-glycero-3-phosphocholine; SLPC: 1-stearoyl-2-linoleoyl-sn-glycero-3-phosphocholine; SONPC: 1-stearoyl-2-(9-oxo)nonanoyl-sn-glycero-3-phosphocholine; SOVPC: l-stearoyl-2-(5-oxovaleroyl)-sn-glycero-3-phosphorylcholine; DNPC: 2-dinonanoyl-sn-glycero-3-phosphocholine; 17:0-22:4 PC-d5: 1-heptadecanoyl-2-docosatetraenoyl-sn-glycero(d5)-3-phosphocholine.
